# Elucidating the molecular programming of a nonlinear nonribosomal peptide synthetase responsible for fungal siderophore biosynthesis

**DOI:** 10.1101/2022.10.10.511241

**Authors:** Matthew Jenner, Yang Hai, Hong H. Nguyen, Munro Passmore, Will Skyrud, Junyong Kim, Neil K. Garg, Wenjun Zhang, Rachel R. Ogorzalek Loo, Yi Tang

## Abstract

Siderophores belonging to the ferrichrome family are essential for the viability of fungal species and play a key role for virulence of numerous pathogenic fungi. Despite their biological significance, our understanding of how these iron-chelating cyclic hexapeptides are assembled by non-ribosomal peptide synthetase (NRPS) assembly lines remains poorly understood, primarily due to the nonlinearity exhibited by the domain architecture. Herein, we report the biochemical characterization of the SidC NRPS, responsible for construction of the intracellular siderophore ferricrocin. *In vitro* reconstitution of purified SidC revealed its ability to produce ferricrocin and its structural variant, ferrichrome. Application of intact protein mass spectrometry uncovered several non-canonical events during peptidyl siderophore biosynthesis, including inter-modular loading of amino acid substrates and an adenylation domain capable of poly-amide bond formation. This work expands the scope of NRPS programming, allows biosynthetic assignment of ferrichrome NRPSs, and sets the stage for reprogramming towards novel hydroxamate scaffolds.

## INTRODUCTION

Iron is an indispensable cofactor for all microbial life. The ability to coordinate and activate molecular oxygen, in addition to optimal redox properties for electron transport, places it central to numerous cellular processes.^1,2^ Equally, high intra-cellular iron concentrations give rise to Fenton and Haber–Weiss reactions, producing reactive oxygen species capable of cell damage.^3^ It is therefore vital that iron homeostasis is carefully managed. Although iron has a high natural abundance, it exists predominantly as Fe^3+^ in aerobic environments and tends to form insoluble ferric hydroxides rendering it inaccessible to microorganisms.^4^ As a result, organisms have evolved complex strategies for iron acquisition and storage. Whilst several mechanisms are known, a common approach employed by bacteria and fungi is the production of low-molecular-weight compounds known as siderophores, which serve as high-affinity iron chelators.^5,6^

In fungi, the majority of siderophore compounds produced belong to the hydroxamate class. This functionality originates from L-ornithine, which is *N^δ^*-hydroxylated and subsequently *N^δ^*-acylated to yield either *N^δ^*-acetyl-*N^δ^*-hydroxy-L-ornithine (AHO) or *N^δ^*-anhydromevalonyl-*N^δ^*-hydroxy-L-ornithine (AMHO).^7^ Typically, siderophores possess three hydroxamate units, producing a hexadentate ligand which promotes formation of a polyhedral Fe^3+^ complex with binding constants in the 10^22^ – 10^32^ range.^8^ The hydroxamate-containing units, AHO and *cis*-*trans*-AMHO, are enzymatically incorporated into chemical scaffolds and define two separate families of hydroxamate siderophores. These include the depsipeptides, typified by fusarinine C (FSC) (**1**),^9^ which utilise either *cis*- or *trans*-AMHO as monomeric units and are excreted primarily to capture ferric iron (**Fig. 1a**).^10^ In contrast, members of the ferrichrome family, such as ferricrocin (**2**) and ferrichrome (**3**), are generally considered to be intracellular and can incorporate AHO or *cis*- / *trans*-AMHO, in combination with other amino acids, and are principally used for iron storage, although not exclusively (**Fig. 1b**, **Supplementary Fig. S1**).^11,12^ Both extra- and intra-cellular siderophores are essential for the survival and virulence of many problematic fungal species, including the opportunistic pathogen *Aspergillus fumigatus* and the rice blast fungus *Magnaporthe oryzae*.^13,14^

**Figure 1.**
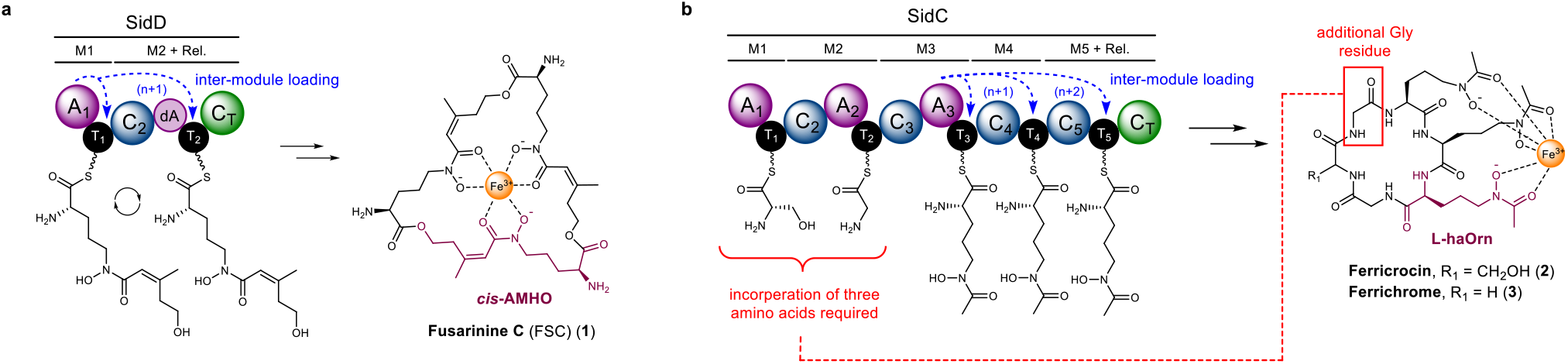
Hydroxamate-containing siderophores produced by fungi and the non-linear NRPSs responsible for their biosynthesis. **a**). Domain organisation of the SidD NRPS responsible for the biosynthesis of fusarinine C (**1**). The A_1_ domain loads *cis*-AMHO units onto the T_1_ and T_2_ domains (highlighted by blue dashed arrows), a requirement due to an inactive A domain (dA) present in module 2. The NRPS acts in an iterative manner to condense three *cis*-AMHO units as a depsipeptide, yielding (**1**) as the final product. **b**). Domain organisation of the SidC NRPS responsible for the biosynthesis of ferricrocin (**2**). The structural variant, ferrichrome (**3**), is also highlighted. It is hypothesised that the A_3_ domain loads L-haOrn units onto the T_4_ and T_5_ domains in a similar manner to SidD (highlighted by blue dashed arrows), as their respective modules lack dedicated A domains. The domains encompassing modules 1 and 2 must incorporate three amino acids [Gly-Ser-Gly] for (**2**) or [Gly-Gly-Gly] for (**3**) (highlighted in red). However, only two A domains are present, indicating unusual nonlinear behaviour of the NRPS. In each case, siderophores are shown in their ferric-bound state, and the hydroxymate-containing monomer unit is highlighted in purple.

Whilst the physiological function of hydroxamate siderophores in fungi is well established, in some cases, the molecular details underpinning their biosynthesis remain poorly understood. Genes encoding for large non-ribosomal peptide synthetase (NRPS) enzymes are known to be responsible for the assembly of peptidyl siderophores.^15–17^ These modular multi-domain enzymes are typically comprised of three domain types: condensation (C), adenylation (A) and thiolation (T). During the biosynthetic process, the peptidyl intermediates are covalently tethered to the T domains via a thioester linkage, afforded by a 4’-phosphopantetheine (Ppant) moiety post-translationally appended to each T domain.^18^ Within a module, the A domain specifically selects and loads an amino acid starter unit (module 1 only) or extender units onto the Ppant thiol of the T domains. This allows the C domain to catalyse amide bond formation between the growing peptidyl intermediate appended to the T domain of the upstream module, and the amino acid extender unit primed on the T domain.^19,20^ Once all cycles of chain elongation are complete, the nascent peptidyl chain is cleaved from the NRPS by either a thioesterase (TE) domain, or more commonly in fungal NRPSs, a C_T_ domain, which catalyse chain-length-specific intramolecular cyclisation to release the final product.^21,22^

In bacterial NRPSs, the colinear relationship between the domain organisation and the final product allows rational assignment of biosynthetic pathways and even prediction of products from sequence data alone.^20^ In contrast, fungal NRPSs typically exhibit highly aberrant domain organisations, making understanding their biosynthetic pathways challenging.^17,23^ A recurring non-canonical feature of fungal NRPSs is modules lacking a functional A domain, implying that the T domain must be loaded by an A domain from a different module. This can be observed in the SidD and Nps6 NRPSs, where module 2 (M2) possesses a truncated A domain (dA) that lacks ~290 amino acids from the N-termini rendering it catalytically inactive (**Fig. 1a**).^24^ Our previous work has demonstrated that the SidD A_1_ domain loads *cis*-AMHO onto both the T_1_ and T_2_ domains, thus circumventing the requirement for an A domain in module 2.^25^ Another distinctive trait is that the length and sequence of the peptide product does not correlate to the number / order of domains in the NRPS, indicative of nonlinear behaviour. This is also exemplified by the SidD NRPS, which incorporates three *cis*-AMHO units yet harbours only 2 modules. Our characterisation of SidD revealed that the NRPS acts in an iterative manner, loading another *cis*-AMHO unit onto the T_1_ domain after formation of the di-*cis*-AMHO species on T_2_, allowing a third *cis*-AMHO residue to be condensed and thereby yielding the fusarinine C (**1**) product (**Fig. 1a**).^25^ Whilst currently limited to the SidD NRPS, these examples of nonlinear behaviour begin to highlight the intricate and highly unusual molecular programming of fungal NRPSs.

The biosynthesis of siderophores belonging to the ferrichrome family have been linked to an evolutionary-related group of NRPSs which exhibit unusual nonlinear domain organisations. An example of one such systems is the SidC NRPS from *Aspergillus nidulans*, which is known to produce ferricrocin (**2**) (**Fig. 1b**).^26^ Variations in the domain architecture of the NRPS exist, some of which give rise to the structurally related ferrichrome (**3**), such as Sib1 from *Schizosaccharomyces pombe* and Sid2 from *Ustilago maydis* (**Supplementary Fig. S1a**).^17,27,28^ The majority of ferrichrome family siderophores are cyclic hexapeptides with the exceptions of the cyclic heptapeptide, tetraglycylferrichrome, and cyclic octapeptide, epichloenin A (**Supplementary Fig. S1b** and **c**).^29,30^ Structurally, they are comprised of three *N^δ^*-acetyl-*N^δ^*-hydroxy-L-ornithine (AHO) residues for Fe^3+^ chelation, and three amino acids forming a variable backbone, of which two residues are either alanine, serine or glycine, and the third a glycine residue.^7^ Similar to SidD, the domain organisation of the SidC NRPS suggests a high degree of nonlinear behaviour. The SidC NRPS possesses only three A domains, yet the siderophore product requires incorporation of six amino acids. Using a linear biosynthetic logic, modules 4 and 5 both lack integrated A domains required for priming the T_4_ and T_5_ domains, presumably with AHO units. However, based on our previous observations of the SidD A_1_ domain, we hypothesise that the SidC A_3_ domain is likely to load the T_4_ and T_5_ domains with AHO units in an inter-modular manner (**Fig. 1b**). Assuming the T_4_ and T_5_ domains are indeed primed by the A_3_ domain, under a linear paradigm this would only permit synthesis of a five-residue peptide product, suggesting further aberrant behaviour to allow incorporation of an additional Gly residue into the backbone of both ferricrocin (**2**) and ferrichrome (**3**) (**Fig. 1b**). It is worth noting that this level of nonlinearity has not been observed before and the exact molecular mechanism cannot be explained by any previous model.

In order to understand the enigmatic programming rules of this particular class of siderophore-producing NRPSs, we have employed a combination of *in vitro* biochemical assays and intact protein mass spectrometry (MS) to interrogate SidC with respect to its ability to produce peptidyl siderophore products. Our results uncover several non-canonical events during peptidyl siderophore biosynthesis, which adds previously unobserved capabilities to NRPS assembly-line enzymology and sets the stage for efforts towards reprogramming the SidC NRPS towards novel chemical scaffolds.

## RESULTS

### Reconstitution of the SidC NRPS and determination of adenylation domain specificity

In the first instance, we elected to examine SidC activity *in vivo* using *Saccharomyces cerevisiae* as a heterologous host. This was conducted to allow production of the associated siderophore(s) and to ascertain whether *S. cerevisiae* would be an appropriate host for recombinant overproduction of SidC for subsequent purification and *in vitro* analysis. To achieve this, the *sidC* NRPS gene from *A. nidulans* FSGC A1145, in addition to *sidA* and *sidL* (required for production of the L-haOrn precursor), were cloned into vectors with distinct selection markers, and transformed into *S. cerevisiae* BJ5464-*npgA* for siderophore production.^31^ Analysis of the small molecule extract from a 3-day culture indicated that ferricrocin (**2**) was produced (**Fig. 2a**, trace i), and large scale cultures allowed purification and isolation of ferricrocin (**2**) for structural elucidation, which was in agreement with previous reports (see **Supplementary Fig. S2 – S4**). Having established that an active form of SidC can be produced in *S. cerevisiae*, recombinant SidC protein was overproduced in *S. cerevisiae* JHY686 as an N-terminal pHis8 fusion protein, and was purified to near-homogeneity using immobilized metal-ion affinity chromatography (IMAC) (**Supplementary Fig. S5**), thereby allowing controlled exposure to substrates / cofactors.^32^ To ensure protein samples were completely in the *holo*-form prior to assays, purified SidC was enzymatically phosphopantetheinylated using the fungal phosphopantetheinyl transferase, NpgA (*A. nidulans*), as described previously.^33^ Following addition of ATP and Mg^2+^ cofactors to recombinant SidC, incubation with L-haOrn alone (synthesised according to literature protocols, **Supplementary Fig. S6 – S8**), or a combination of L-haOrn + L-Ser, yielded no detectable products (**Fig. 2a**, trace ii and iii). However, the combination of L-haOrn + Gly + L-Ser resulted in the production of ferricrocin (**2**) (**Fig. 2a**, trace v). Interestingly, incubation with L-haOrn + Gly resulted in formation of a species consistent with ferrichrome (**3**) (**Fig. 2a**, trace iv), which was confirmed by comparison to a chemical standard (**Fig. 2a**, trace vii). These observations suggest that the SidC NRPS is capable of producing both ferricrocin (**2**) and ferrichrome (**3**) depending upon the availability of amino acid substrates, yet appears to produce exclusively ferricrocin (**2**) in the native host, probably due to the abundance of L-Ser.

**Figure 2.**
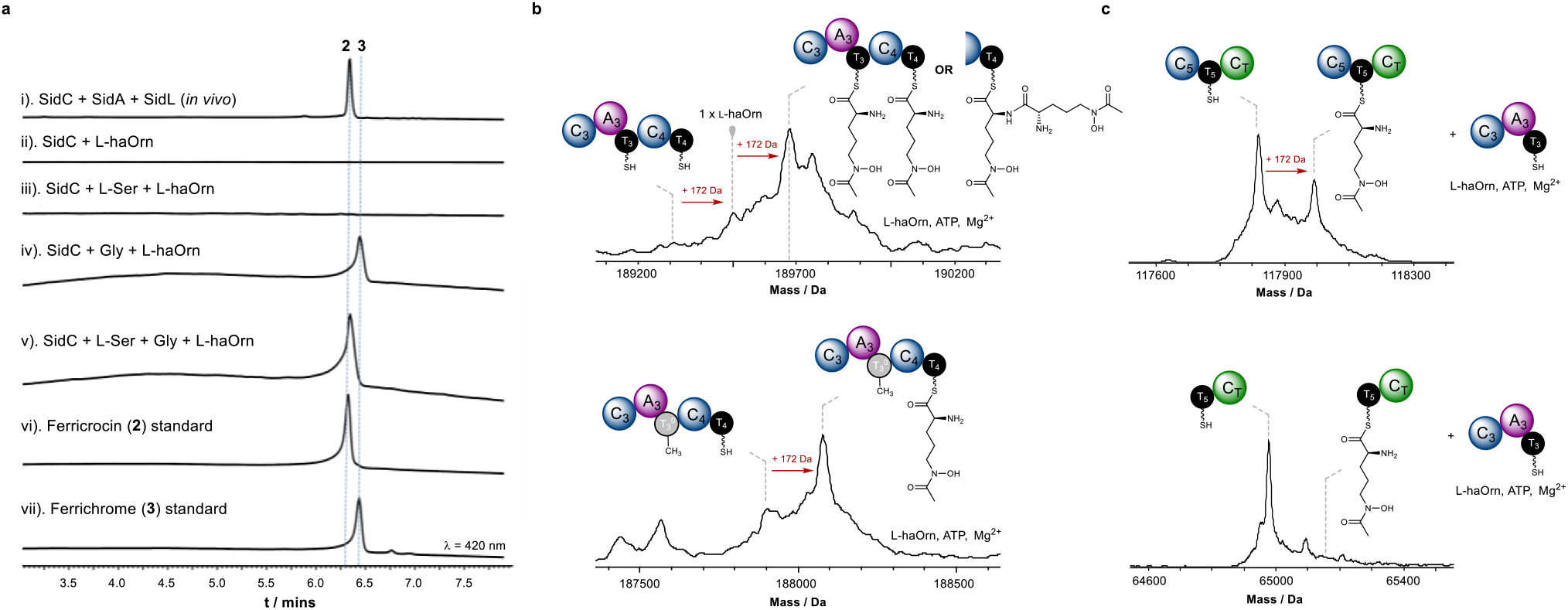
Reconstitution of SidC NRPS and inter-modular loading of L-haOrn residues by the A_3_ domain. **a**). HPLC traces monitored at 420 nm for the following: i). production of **2** via heterologous expression of *sidC, sidA* and *sidL* in *S. cerevisiae* JHY686; ii) - v). *in vitro* enzymatic reactions of SidC in the presence of L-haOrn, +/− L-Ser and +/− Gly; vi – vii). authentic standards of **2** and **3**. Presented with either a cellular pool of amino acids, or L-haOrn + L-Ser + Gly *in vitro*, SidC produces **2** exclusively. However, when provided with L-haOrn + Gly *in vitro*, SidC produces solely **3**. **b**). Deconvoluted intact protein mass spectra of *holo*-SidC C_3_A_3_T_3_C_4_T_4_ (*top*) following incubation with L-haOrn, ATP and Mg^2+^, showing loading of either: x2 L-haOrn units onto the T_3_ and T_4_ domains, or a condensed di-L-haOrn species on the T_4_ domain. *holo*-SidC C_3_A_3_T_3_^0^C_4_T_4_ (*bottom*) following incubation with L-haOrn, ATP and Mg^2+^, showing loading of a single L-haOrn unit onto the T_4_ domain. The S3151A mutation in the T_3_ domain means it is unable to be modified with a Ppant moiety, thus preventing loading of L-haOrn. **c**). Deconvoluted intact protein mass spectra of *holo*-SidC C_5_T_5_C_T_ (*top*) and *holo*-SidC T_5_C_T_ (*bottom*) following incubation with *holo*-SidC C_3_A_3_T_3_, L-haOrn, ATP and Mg^2+^. Loading of L-haOrn is only observed when the N-terminal C domain of each construct is present. Mass shifts corresponding to biosynthetic steps are highlighted with red arrows, and proposed intermediates are displayed. Exact measured and observed masses are detailed in **Table S2**.

Our initial biosynthetic model hypothesised that each of the three A domains within SidC are responsible for activation and loading of L-Ser, Gly and L-haOrn (**Fig. 1b**). In bacteria, bioinformatic analysis allows accurate prediction of the substrate specificity for A domains, primarily based on highly conserved amino acid motifs within the enzyme active site. However, this approach is not possible for fungal A domains, largely due to limited sequence and structural information. Unable to predict the substrate specificity of the SidC A_1_, A_2_ and A_3_ domains using bioinformatics, excised constructs of SidC A_1_, A_2_ and A_3_ were cloned, overproduced in *E. coli* and purified to homogeneity for determination of substrate specificity. Using an ATP/PPi exchange assay to measure the reverse reaction of the adenylation step, the SidC A_1_ domain showed activation of L-Ala and L-Ser, with a clear preference towards the latter (**Supplementary Fig. S9,** *top*). Interestingly, the SidC A_2_ domain exhibited no detectable activity when subjected to the ATP/PPi exchange assay, suggesting that PPi may not formed during the activation step. However, activity was observed for SidC A_2_ using the hydroxylamine release assay, which indicates formation of the corresponding aminoacyl-adenylate, and allowed a substrate specificity profile to be obtained revealing Gly as the preferred substrate (**Supplementary Fig. S9,** *middle*). The discrepancy between these two assays indicates that the A_2_-catalysed adenylation reaction is not reversible.^34,35^ Activity for the SidC A_3_ domain was obtained using the ATP/PPi exchange assay and displayed a clear preference towards L-haOrn (**Supplementary Fig. S9,** *bottom*). The SidC A_1_ and A_2_ assays were also reconstituted with their cognate T_1_ and T_2_ domains to examine the aminoacyl transfer step. Intact protein mass spectrometry (MS) analysis showed mass shifts loading of L-Ser and Gly onto the respective T domains (**Supplementary Fig. S10**).

### The SidC A_3_ domain catalyses *intra*- and *inter*-modular loading of L-haOrn

The biosynthesis of ferricrocin (**2**) and ferrichrome (**3**) both require installation of three L-haOrn residues, yet the SidC NRPS possesses only a single A domain capable of activating L-haOrn; the A_3_ domain situated in module 3. Furthermore, modules 4 and 5 lack integrated A domains for loading amino acid units to their cognate T domains (**Fig. 1b**). Based on our previous observations in the SidD NRPS, we postulated that the SidC A_3_ domain may be capable of loading L-haOrn units onto the T_4_ and T_5_ domains, in addition to its cognate T_3_ domain. This would result in the T_3_, T_4_ and T_5_ domains being charged with L-haOrn units, which can then be condensed together by the sequential activity of the C_4_ and C_5_ domains to yield the tri-L-haOrn motif. An alternative model would involve iterative activity of module 3 to generate tri-L-haOrn appended to the T_3_ domain, however this would render the C_4_T_4_C_5_T_5_ region redundant for the biosynthesis and seemed less likely.

To investigate this aspect, a SidC C_3_A_3_T_3_ tri-domain construct was cloned, overproduced and purified to examine the covalently tethered intermediates loaded onto the T_3_ domain. Following conversion to its *holo* form and subsequent incubation with ATP, Mg^2+^ and L-haOrn (15 min), intact protein MS analysis of C_3_A_3_T_3_ revealed loading of a single L-haOrn unit, indicated by a +172 Da mass shift relative to the mass of *holo*-C_3_A_3_T_3_ (**Supplementary Fig. S11**). This highlighted that the standalone SidC C_3_A_3_T_3_ tri-domain is only capable of loading a single L-haOrn unit onto its cognate T_3_ domain and ruled out the possibility of iterative loading. We next generated a SidC C_3_A_3_T_3_C_4_T_4_ penta-domain construct to examine the ability of the A_3_ domain to catalyse loading of L-haOrn onto the T_4_ domain. Under the same conditions, two sequential +172 Da mass shifts were observed in the mass spectrum, congruent with loading of two L-haOrn units (**Fig. 2b**, *top*). The measured masses were consistent with either two L-haOrn units loaded in an uncondensed form onto the T_3_ and T_4_ domains, or in a condensed di-L-haOrn species on the T_4_ domain. To validate these observations, a mutant of the SidC C_3_A_3_T_3_C_4_T_4_ construct was produced, where the Ser residue that serves as the Ppant group attachment site was mutated to Ala (S3140A, designated as T_3_^0^), allowing only the T_4_ domain to be converted to its *holo* form. When subjected to the loading assay, the SidC C_3_A_3_T_3_^0^C_4_T_4_ protein was able to activate and transfer an L-haOrn unit onto the T_4_ domain, indicated by a single +172 Da mass shift in the intact protein mass spectrum (**Fig. 2b,** *bottom*).

Inter-modular loading of T_4_ by the A_3_ domain in the SidC NRPS is reminiscent of behaviour observed for the SidD NRPS, where the A_1_ domain is able to prime T_2_, situated in the downstream module. In both of these instances, the A domain is interacting with a T domain situated one module downstream. However, in the SidC NRPS, we hypothesised that the A_3_ domain is capable of loading the T_5_ domain, situated two modules downstream. To probe this, we conducted a bimolecular assay between SidC C_3_A_3_T_3_ and SidC C_5_T_5_C_T_ in the presence of ATP, Mg^2+^ and L-haOrn. This resulted in ~35 % of L-haOrn loading onto the T_5_ domain after a 60 min incubation (**Fig. 2c**, *top*), and indicated that the A_3_ domain is capable of loading the T_5_ domain, situated two modules downstream. An equivalent experiment using a C_4_T_4_ construct yielded comparable levels of L-haOrn loading, serving as a control measurement, and also highlighting the reduced efficiency when domains are not covalent tethered in megasynth(et)ases (**Supplementary Fig. S12**). Interestingly, spectra obtained from this experiment also gave rise to a small peak congruent with a di-L-haOrn species. The relatively small amount of this condensed species relative to the mono-L-haOrn suggests that the C_3_ domain does not preferentially condense L-haOrn units together, indicating that the species in **Fig. 2b** (*top*) is likely two uncondensed L-haOrn units. Analogous assays using the SidC T_5_C_T_ didomain and SidC T_4_ domain (i.e. without the N-terminal C domain), resulted in no detectable L-haOrn loading (**Fig. 2c**, *bottom* and **Supplementary Fig. S12**), implying that the N-terminal C domains facilitate the loading reaction. These results suggest two architectural models for non-linear L-haOrn loading by the A_3_ domain. One possibility is that intra-chenar loading of L-haOrn is promoted by a 3-dimensional arrangement of the SidC NRPS that enables proximity of the A_3_ domain to the T_4_ and T_5_ domains (**Supplementary Fig. S13**). Here, the presence of the C_4_ and C_5_ domains may be essential to provide an interaction ‘platform’ for the T_4_ and T_5_ domains to access the A_3_ domain. A second possibility involves inter-chenar communication between two SidC proteins, allowing the A_3_ domain to load L-haOrn onto the T_4_ and T_5_ domain *in trans*, whilst loading the T_3_ domain conventionally (**Supplementary Fig. S13**).

### SidC C_2_A_2_T_2_ tri-domain catalyses non-canonical loading of Gly residues

We next turned our attention to the biosynthetic steps required for the formation of the tripeptide backbone. This region is the sole structural difference between ferricrocin (**2**) and ferrichrome (**3**), possessing [Gly]-[L-Ser]-[Gly] and [Gly]-[Gly]-[Gly] motifs, respectively. The SidC A domain specificity assays determined that the A_1_ domain preferentially activates L-Ser, and that the A_2_ domain only activates Gly (**Supplementary Fig. S9** and **S10**). Based on these observations, linear assembly of the amino acid units would yield a T_2_-[Gly]-[L-Ser]-NH_2_ species, requiring a second Gly to be non-canonically condensed onto the amine of L-Ser to yield the T_2_-[Gly]-[L-Ser]-[Gly]-NH_2_ tripeptide intermediate necessary for ferricrocin (**2**) production. However, for ferrichrome (**3**), two scenarios seemed plausible: i). the A_1_ domain would instead load Gly onto the T_1_ domain (note, some activity towards Gly observed in specificity assays (**Supplementary Fig. S9**)), allowing a T_2_-[Gly]-[Gly]-NH_2_ species to be formed, followed by non-canonical condensation of a third Gly to yield the T_2_-[Gly]-[Gly]-[Gly]-NH_2_ intermediate. ii). the A_1_ and T_1_ domains are not utilised, leaving the A_2_ domain to generate a T_2_-[Gly]-NH_2_ species, which must undergo two sequential non-canonical condensation events to yield the T_2_-[Gly]-[Gly]-[Gly]-NH_2_ intermediate.

In order to unpick these biosynthetic steps, we first produced a SidC(ΔA_1_T_1_) construct to examine whether the domains of module 1 are essential for siderophore production. Using *S. cerevisiae* as a heterologous host, SidC (ΔA_1_T_1_) was observed to produce ferrichrome (**3**) exclusively, with no ferricrocin (**2**) detected (**Fig. 3a**, trace i), which was the product of full-length SidC under the same culture conditions (**Fig. 2a**). Purification of the recombinant SidC(ΔA_1_T_1_) protein allowed controlled exposure to amino acid substrates. Here, SidC(ΔA_1_T_1_) produced ferrichrome (**3**) exclusively, provided that both Gly and L-haOrn were present (**Fig. 3a**, traces ii – iii). The inclusion of L-Ser did not promote ferricrocin (**2**) production (**Fig. 3a**, trace ii), and omission of L-haOrn resulted in no detectable products (**Fig. 3a**, trace iv). These data indicated that the A_1_T_1_ domains are essential for ferricrocin (**2**) production, but are not required for ferrichrome (**3**) formation. Furthermore, these data indicated that during ferrichrome (**3**) biosynthesis, neither A_1_ or T_1_ domain participate in the recruitment of the additional Gly residue. This left the intriguing possibility that module 2 alone (i.e. C_2_A_2_T_2_) could be responsible for constructing the T_2_-[Gly]-[Gly]-[Gly]-NH_2_ intermediate required for ferrichrome (**3**) biosynthesis.

**Figure 3.**
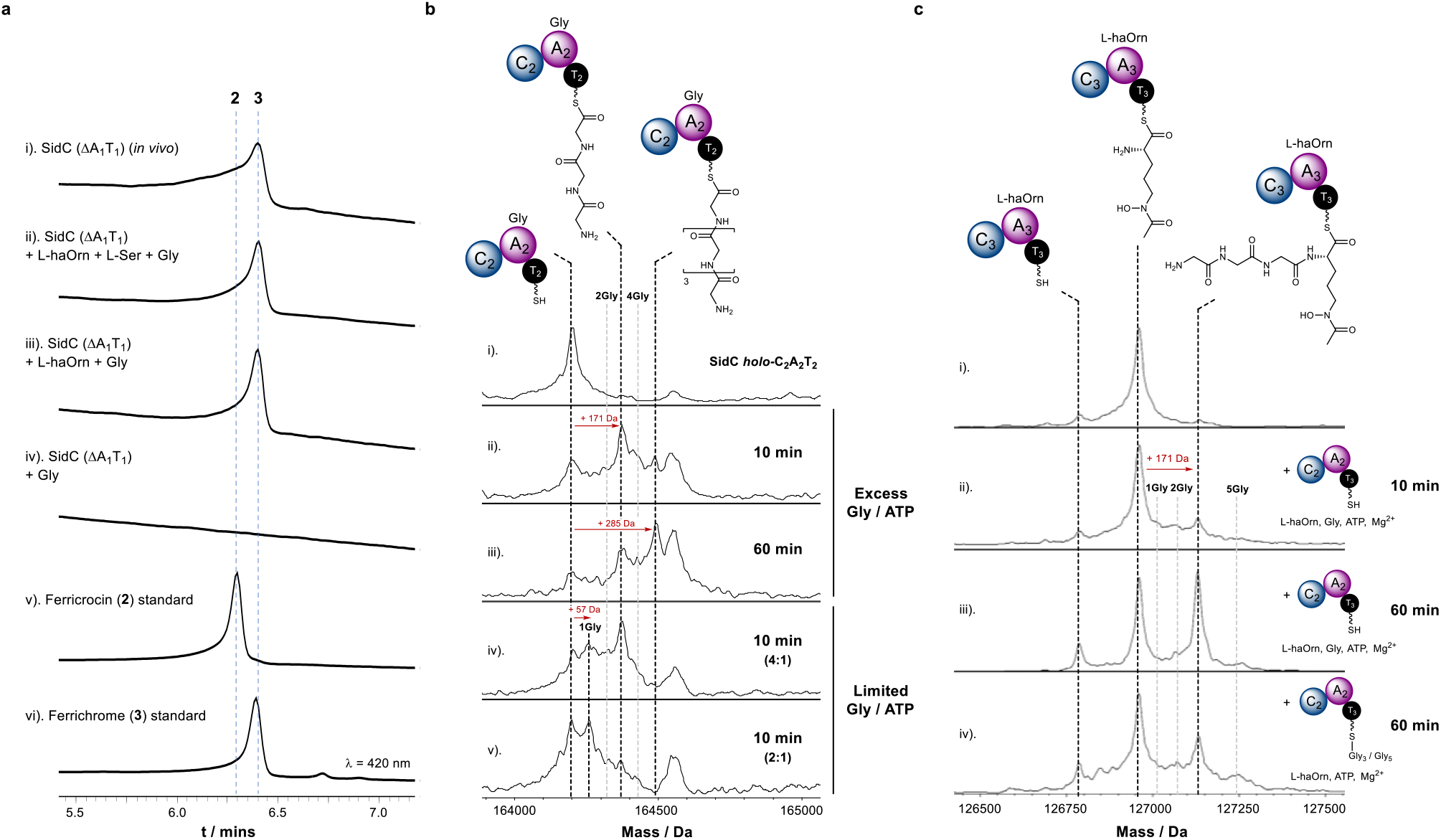
Iterative loading of Gly residues by the A_2_ domain and chain-length control by the C_3_ domain. **a**). HPLC traces monitored at 420 nm for the following: i). production of **3** via heterologous expression of *sidC* (ΔA_1_T_1_), *sidA* and *sidL* in *S. cerevisiae* JHY686; ii - iv). *in vitro* enzymatic reaction of SidC (ΔA_1_T_1_) in the presence of Gly, +/− L-Ser and +/− L-haOrn; v - vi). authentic standards of **2** and **3**. **b**). Stacked deconvoluted intact protein mass spectra of *holo*-SidC C_2_A_2_T_2_ in the presence of Gly, ATP and Mg^2+^. Assays conducted with excess Gly / ATP are shown in spectra ii). and iii). at 10 min and 60 min time intervals, and with limited concentrations of Gly / ATP in spectra iv). and v). after a 10 min incubation. Mass shifts corresponding to mono-/poly-Gly species are highlighted with red arrows, and proposed intermediates are displayed. **c**). Stacked deconvoluted intact protein mass spectra of L-haOrn-SidC C_3_A_3_T_3_ following incubation with *holo*-SidC C_2_A_2_T_2_, Gly, ATP and Mg^2+^ following 10 min and 60 min incubation periods. Spectrum i. shows L-haOrn SidC C_3_A_3_T_3_ alone; spectra ii. and iii. show increasing production of the condensed product, L-haOrn-Gly_3_-SidC C_3_A_3_T_3_ over time. Only the Gly_3_ condensed product is observed, not Gly_1_ or Gly_2_, suggesting that this is not a stepwise process. Instead, Gly_3_ must be formed on SidC C_2_A_2_T_2_ before the SidC C_3_ domain will catalyse the condensation reaction.Spectrum iv. shows an experiment where a 60 min pre-incubation of *holo*-SidC-C_2_A_2_T_2_ with Gly, ATP and Mg^2+^ was conducted to allow formation of Gly_3_/Gly_5_-SidC C_2_A_2_T_2_ (see **Fig. 3b**, spectrum iii.), before addition of L-haOrn-SidC C_3_A_3_T_3_. Only the Gly_3_ condensed product is observed, not Gly_5_, indicating that the SidC C_3_ domain selectively condenses the Gly_3_-SidC C_2_A_2_T_2_ species with L-haOrn only. Mass shifts corresponding to biosynthetic species are highlighted with red arrows, and proposed intermediates are displayed. Exact measured and observed masses are detailed in **Table S2** and **S3**.

To explore this prospect, we cloned an MBP-SidC C_2_A_2_T_2_ fusion construct, which was overexpressed in *E. coli* as a soluble protein and purified to homogeneity for *in vitro* studies. Using our established intact protein MS approach, incubation of SidC C_2_A_2_T_2_ with excess Gly, ATP and Mg^2+^ for 10 mins led to emergence of two new peaks which were + 171 Da and + 285 Da relative to the *holo*-C_2_A_2_T_2_ species, mass shifts which correspond to condensed tri- and penta-Gly chains, respectively, attached to the T_2_ domain (**Fig. 3b**, spectra i – ii). Upon extension of the incubation period to 60 min, the relative abundance of the penta-Gly species increased as the tri-Gly decreased, suggesting that the tri-Gly intermediate is a precursor to the penta-Gly chain (**Fig. 3b**, spectrum iii). In the presence of excess Gly and ATP conversion to the tri-Gly and penta-Gly intermediates was fast, preventing observation of early intermediates of the poly-Gly chain. Therefore, assays were repeated with reduced relative concentrations of 4:1 and 2:1 (Gly/ATP:protein) to capture early-stage intermediates. At a 4:1 ratio, a peak at + 57 Da relative to the *holo*-C_2_A_2_T_2_ species was observed, which correlates to a single Gly unit loaded onto the T_2_ domain, in addition to the previously observed tri-Gly species (**Fig. 3b**, spectrum iv). Switching to a 2:1 ratio yielded predominantly the mono-Gly species, with low levels of di- and tri-Gly intermediates observable in the spectrum (**Fig. 3b**, spectrum v).

These observations indicated that the domains of module 2 alone are capable of constructing Ppant-bound poly-Gly chains, appearing to favour the formation of tri- and penta-Gly species. While loading of a single Gly unit onto the T_2_ domain is likely catalysed by the A_2_ domain in a canonical fashion, the mechanism by which the remaining two Gly units are condensed onto the free amine remained elusive. The use of an isolated C_2_A_2_T_2_ construct in our experiments, meant it was possible that inter-chenar communication between individual C_2_A_2_T_2_ proteins could allow the C_2_ domain to catalyse condensation reactions between Gly chains, assuming that the T_2_ domain is able to act as an aminoacyl donor and acceptor (i.e. C_2_A_2_T_2_-[Gly]-NH_2_ + C_2_A_2_T_2_-[Gly]-NH_2_ → C_2_A_2_T_2_-SH + C_2_A_2_T_2_-[Gly]-[Gly]-NH_2_). To test this possibility, the catalytically essential active site His_569_ residue of the C_2_ domain was mutated to Ala to produce a catalytically inactive C_2_ domain, denoted as C_2_^0^. Upon incubation with Gly and co-factors, the SidC C_2_^0^A_2_T_2_ protein was able to generate poly-Gly chains in a near-identical manner to that of the wild-type construct (**Supplementary Fig. S14**). We attempted to produce a SidC A_2_T_2_ construct, however, in the absence of the N-terminal C_2_ domain the resulting protein was highly unstable and degraded quickly, suggesting that the C domain plays an important structural role within the module.

Taken together, our data strongly indicate that the SidC A_2_ domain is responsible for catalysing both canonical loading of a single Gly unit onto the Ppant thiol of the T_2_ domain, and subsequent amide bond formation steps to generate the tri-Gly intermediate required for ferrichrome (**3**) biosynthesis. Whilst two non-canonical amide bond-forming steps are required for ferrichrome (**3**) biosynthesis, this is only required once for ferricrocin (**2**) biosynthesis, adding a Gly to the free amine of the T_2_-[Gly]-[L-Ser]-NH_2_ species. Poly-amide bond formation catalysed by an A domain is unprecedented within the context of a multi-modular NRPS, making this system particularly interesting. However, similar activity has been observed for a standalone A domain during the biosynthesis of streptothricin. Here, following canonical loading of a L-β-lysine residue onto a T domain, a separately encoded adenylation domain, ORF19, generates poly-L-β-lysine chains via amide bond formation with the free ε-NH_2_ group (**Supplementary Fig. S15**).^36^ Interestingly, the SidC A_2_ domain differs from ORF19, in that the amide bond is formed using the α-NH_2_ group and the domain is found integrated into the NRPS. It is worth noting that stand-alone adenylation domains have also been observed to catalyse amide bond formation in the biosynthesis of pacidiamycin (PacU), coumermycin A_1_ (CouL) and novobiocic acid (NovL) (**Supplementary Fig. S15**).^37–39^ However, these examples involve single condensation events, not formation of poly-amino acid chains as observed for the ORF19 and SidC A_2_ domains.

### The SidC C_3_ domain is a chain-length gatekeeper

Our biochemical investigations of SidC C_2_A_2_T_2_ demonstrated its ability to generate poly-Gly chains of up five residues in length (**Fig. 3b**). However, the biosynthetic products of SidC, ferricrocin (**2**) and ferrichrome (**3**), both require the T_2_-tethered intermediate to be three residues in length: T_2_-[Gly]-[L-Ser]-[Gly]-NH_2_ and T_2_-[Gly]-[Gly]-[Gly]-NH_2_, respectively. Therefore, in order to maintain biosynthetic fidelity, we postulated that the SidC C_3_ domain imposes a selective requirement for three-residue chains appended to the T_2_ domain in order to catalyse condensation with the first L-haOrn unit, effectively acting as a gatekeeper. To explore this hypothesis, we incubated *holo*-SidC C_2_A_2_T_2_ with *holo*-SidC C_3_A_3_T_3_ in the presence of Gly, L-haOrn, ATP and Mg^2+^, and monitored the SidC C_3_A_3_T_3_ protein using intact protein MS at several time points. After 10 min, a new peak at + 171 Da from the L-haOrn-C_3_A_3_T_3_ species had emerged, indicating condensation of a tri-Gly unit onto the L-haOrn (**Fig. 3c**, spectra i and ii), with the intensity of this species increasing over a 60 min period (**Fig. 3c**, spectrum iii). The absence of signals corresponding to mono- (+ 57 Da) or di-Gly (+ 114 Da) species condensed with L-haOrn-C_3_A_3_T_3_ during the time-course strongly indicated that the entire T_2_-[Gly]-[Gly]-[Gly] intermediate is condensed with L-haOrn, rather than sequential addition of Gly residues.

To examine whether the SidC C_3_ domain can discriminate between tri-, tetra- and penta-Gly intermediates, we pre-incubated *holo*-SidC C_2_A_2_T_2_ with Gly, ATP and Mg^2+^ for 60 min to generate a mixture of poly-Gly chain lengths, represented by **Fig. 3c**, spectrum iii. The remaining Gly in the reaction was then removed by multiple cycles of ultrafiltration, before addition to *holo*-SidC C_3_A_3_T_3_ in the presence of L-haOrn, ATP and Mg^2+^ for 60 min. Subsequent intact protein MS analysis revealed only the tri-Gly species condensed with T_3_-tethered L-haOrn (**Fig. 3c**, spectrum iv), suggesting that the SidC C_3_ domain possesses strict selectivity for poly-amino acid chain lengths where n = 3, thereby acting as a critical checkpoint during the biosynthesis.

### A biosynthetic model for siderophore production by the SidC NRPS

Taken together, our data allows proposal of a rational biosynthetic model for the construction of ferricrocin (**2**) and ferrichrome (**3**) by the SidC NRPS (**Fig. 4**). In ferricrocin (**2**) biosynthesis, the process is initiated by the A_1_ domain loading a L-Ser residue onto the T_1_ domain, which is subsequently condensed with a Gly residue tethered to the downstream T_2_ domain by the C_2_ domain, as a result of A_2_ domain loading, yielding the T_2_-[Gly]-[L-Ser]-NH_2_ intermediate (**Fig. 4a**). This initial step is not required for ferrichrome (**3**) biosynthesis, which commences with A_2_ domain-catalysed loading of a single Gly residue onto the T_2_ domain, and is then condensed with a second Gly residue, catalysed by the amide bond-forming capabilities of the A_2_ domain, yielding a T_2_-[Gly]-[Gly]-NH_2_ intermediate (**Fig. 4b**). Both T_2_-tethered dipeptide intermediates during ferricrocin (**2**) and ferrichrome (**3**) biosynthesis then undergo addition of a Gly residue to the free NH_2_ group, catalysed by the A_2_ domain, producing T_2_-[Gly]-[L-Ser]-[Gly]-NH_2_ and T_2_-[Gly]-[Gly]-[Gly]-NH_2_ intermediates. Whilst the A_2_ domain is capable of adding further Gly residues to extend the peptidyl chain over time (**Fig. 3b**), the nascent tripeptide intermediates are rapidly and selectively condensed with the T_3_-tethered L-haOrn species by the C_3_ domain.

**Figure 4.**
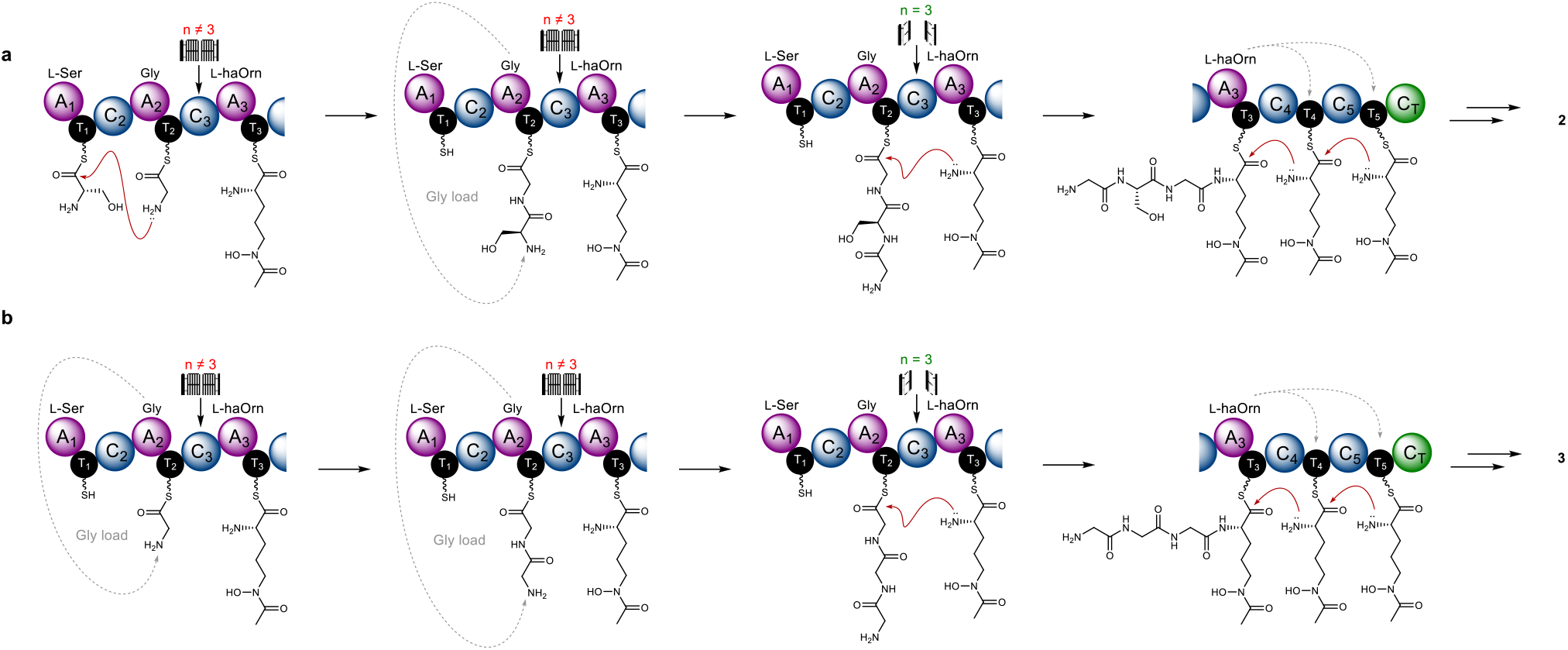
Proposed biosynthetic models for SidC-catalysed formation of ferricrocin and ferrichrome. **a**). Ferricrocin (**2**) biosynthesis commences with condensation between L-Ser and Gly, which is catalysed by the SidC C_2_ domain forming an (L-Ser)-Gly dipeptide (n = 2) tethered to the SidC T_2_ domain. Non-canonical ligation of a Gly unit onto the amine of L-Ser produces a Gly-(L-Ser)-Gly tripeptide (n = 3), which can undergo condensation with L-haOrn catalysed by the chain-length selective SidC C_3_ domain. The SidC A_3_ domain loads L-haOrn onto SidC T_4_ and T_5_ domains allowing a succession of condensation events to generate a Gly-(L-Ser)-Gly-(L-haOrn)_3_ hexapeptide intermediate bound to the SidC T_5_ domain. Chain release is catalysed by the C-terminal C_T_ domain to yield the biosynthetic product **9**. **b**). Ferrichrome (**3**) biosynthesis can occur in the absence of L-Ser, where canonical loading of Gly onto the T_2_ domain is followed by two successive rounds of non-canonical Gly ligation to yield a Gly_3_ species (n = 3) tethered to the T_2_ domain. The remaining steps are identical to the biosynthesis of **2**, to yield the biosynthetic product **3**.

## DISCUSSION

Ferrichrome NRPSs are found in the vast majority of Ascomycetes, providing the biosynthetic machinery for siderophore production. Despite producing near-identical products, ferricrocin (**2**) and ferrichrome (**3**), substantial differences in the NRPS domain architecture exist within the NRPS family (Types I – V, **Fig. 5**). Phylogenetic work has suggested that ferrichrome NRPSs originate from an ancestral colinear hexamodule NRPS, created by adjacent duplication of complete NRPS modules resulting in two lineages: NSP2 and NSP1 / SidC. The recently reported Sid1 NRPS responsible for AS2488059 biosynthesis, a related ferrichrome siderophore, is a contender for this ancestral gene (**Supplementary Fig. S16**).^40,41^ Subsequent losses of individual domains or complete modules from this ancestral gene provides a rational explanation for the diversity of domain architectures.^17^ However, all combinations of the NRPS give rise to unusual non-linear domain organisations, which cannot be reconciled with standard biosynthetic logic of NRPSs. Plausible biosynthetic proposals linking the domain organisation to the peptidyl product require inter-modular loading of amino acid substrates by A domains up to n+2 modules downstream (blue arrows), and / or A domains capable of creating (poly)amide chains on the same T domain (red arrows). Our study of the SidC NRPS highlights that both activities are possible in NRPSs and allow evidence-based biosynthetic proposals for all variations of the ferrichrome NRPS (**Fig. 5**).

**Figure 5.**
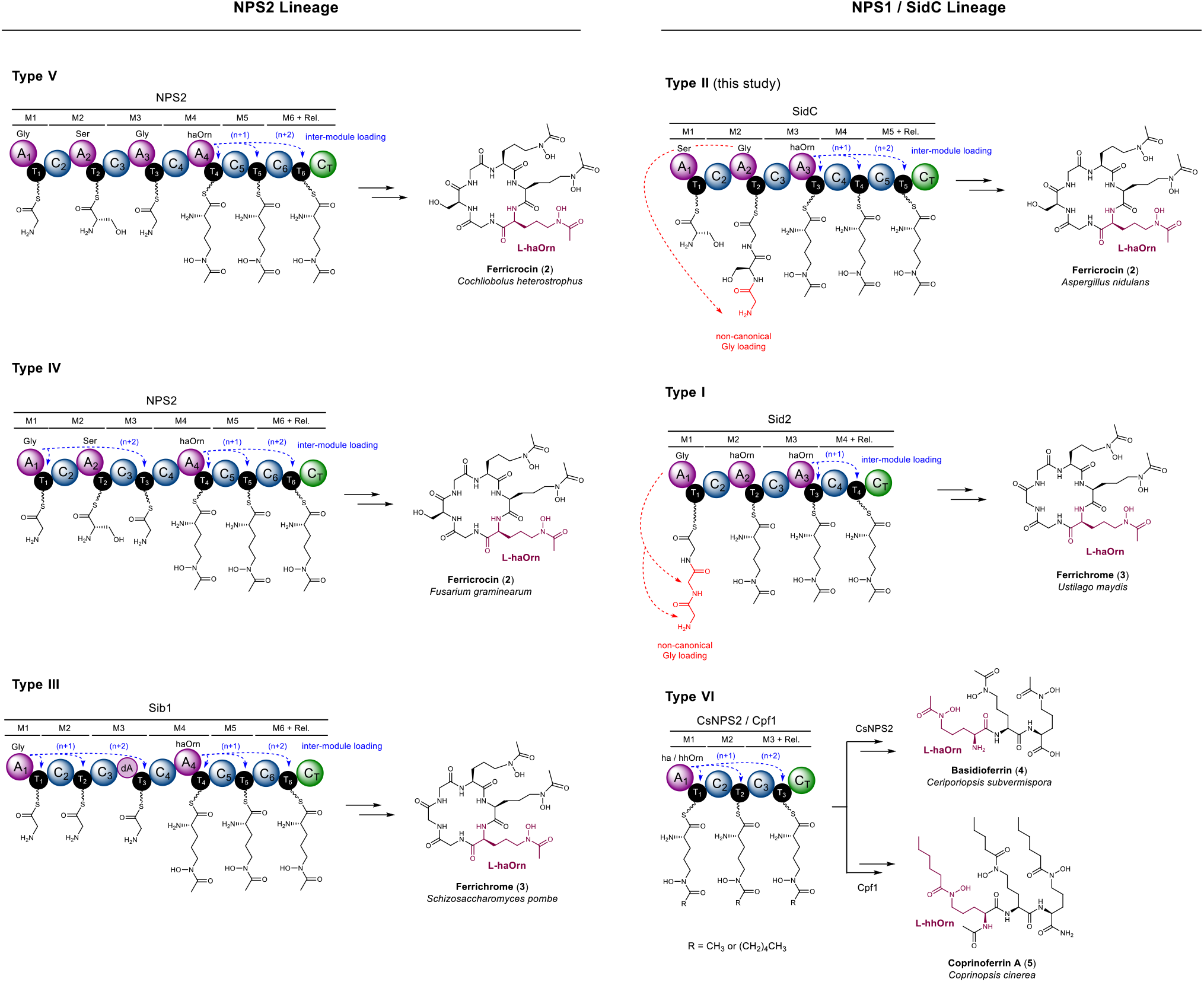
Biosynthetic schemes for the six modular architectures of ferrichrome class of NRPSs. The new programming rules allow the biosynthetic assignment of other architectures of ferrichrome-family synthetase NRPSs. Examples include: Sid2 (ferrichrome), *U. maydis*;^44^ SidC (ferricrocin), *A. nidulans*;^26^ Sib1 (ferrichrome), *S. pombe*;^15,45^ NPS2 (ferricrocin), *F. graminearum*;^46^ NPS2 (ferricrocin), *C. heterostrophus*;^47^ CsNPS2 (basidioferrin), *C. subvermispora*;^42^ and Cpf1 (coprinoferrin A), *C. cinerea*.^43^ The inter-module loading events, either (n+1) or (n+2), are highlighted in blue, and non-canonical loading of Gly residues is highlighted in red. In each case, siderophores are shown in their desferric state, and the hydroxymate-containing monomer unit is highlighted in purple. The lineage classification and Type I – VI groupings are based on previous phylogenetic analyses of ferrichrome synthetase NRPSs conducted by Bushley et al.^17^

Members of the NPS2 lineage (Types III, IV and V) all possess the correct number of C domains required for the number of amide bonds formed in the peptidyl product. However, loss or degeneration of A domains requires inter-module loading of T domains with amino acid substrates. This is observed in the tri-Orn region, as characterised for SidC, and in the tripeptide region for Type III and IV. In contrast, the Type I and II members of the NPS1 / SidC lineage both require an amide bond-forming A domain to compensate for the lack of C domains in the NRPS, in addition to inter-module loading capabilities in the tri-Orn region. The CsNPS2 and Cpf1 NRPSs (Type VI) are the most truncated variation and appears to have lost much of the N-terminus, leaving the just the tri-Orn region that requires inter-module loading capabilities. Recent assignments of the CsNPS2 and Cpf1 products as basidioferrin (**4**) and coprinoferrin (**5**) revealed a structures comprised of three condensed L-haOrn units for basidioferrin (**4**)^42^, and three L-hydroxyhexanoyl ornithine (L-hhOrn) units for coprinoferrin (**5**).^43^ In the latter, this suggests that the Cpf1 A_1_ domain (equivalent to A_3_ in SidC and Sid2) has evolved specificity towards a larger hydroxamate substrate, whilst retaining the ability to conduct inter-module loading (**Fig. 5**).

Taken together, our observations highlight the impressive evolutionary changes employed by fungal NRPSs to improve atom economy and increase structural diversity in their biosynthetic assembly-lines. Our improved understanding of the biosynthetic rules has set the stage for manipulating and recombining these pathways towards novel hydroxamate-containing scaffolds.

## Acknowledgements

This work was supported by a NIH 1R35GM118056 grant to Y. T., and a Life Sciences Research Foundation Fellowship to Y. H. sponsored by the Mark Foundation for Cancer Research. M. J. is the recipient of a BBSRC Discovery Fellowship (BB/R01212/1), and M. P. is supported by a Midlands Integrative Bioscience Doctoral Training Partnership studentship (BB/M01116X/1). The Bruker MaXis II instrument used in this study was funded by the BBSRC (BB/M017982/1). R. R. O. L. and H. H. N. acknowledge funding from the U.S. Department of Energy (DOE) Office of Science (BER) contract DE-FC-02-661 02ER63421 (PI Yeates; UCLA/DOE Institute for Genomics and Proteomics). W. Z. and W. S. acknowledge funding from NIH DP2AT009148 grant. N. K. G. acknowledges shared instrumentation grants from the NSF (CHE-1048804) and the National Center for Research Resources (S10RR025631).

