## Supplementary Information for "Elucidating the molecular programming of a nonlinear nonribosomal peptide synthetase responsible for fungal siderophore biosynthesis"

### 1. Supplementary Figures

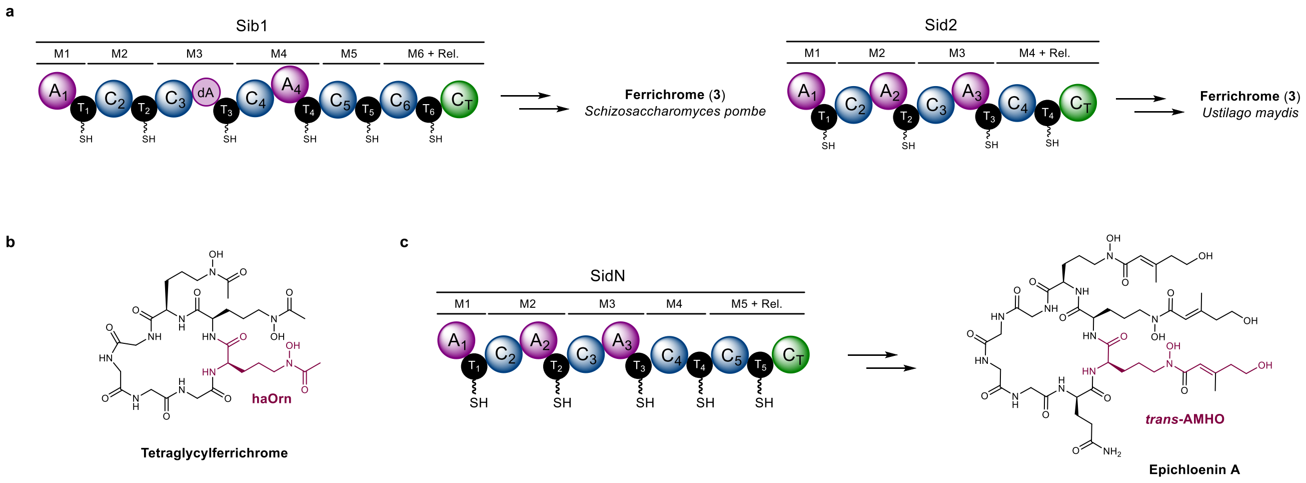


**Figure S1: Variations on ferrichrome family siderophores from fungi and their biosynthetic origins.**
**a**). Domain organisation of the Sib1 and Sid2 NRPSs responsible for production of ferrichrome (**3**). Both NRPSs have subtle variations in their domain organisation to each other, yet produce the same peptidyl product. **b**). Structure of tetraglycylferrichrome. **c**). Structure of epichloenin A and domain organisation of the biosynthetic NRPS, SidN.

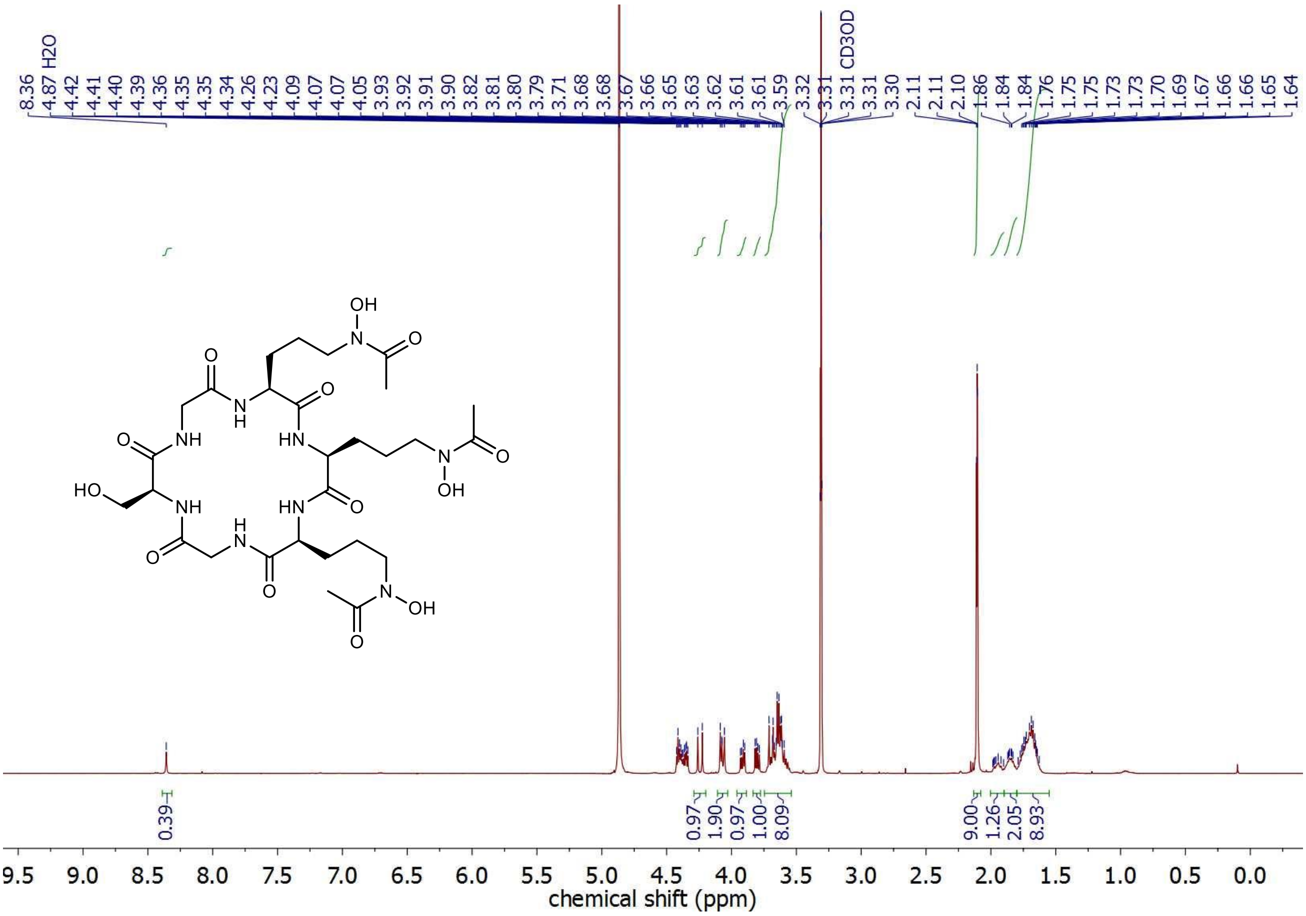


**Fig. S2: [1]H NMR spectrum of desferri-ferricrocin in $CD_3OD$.**

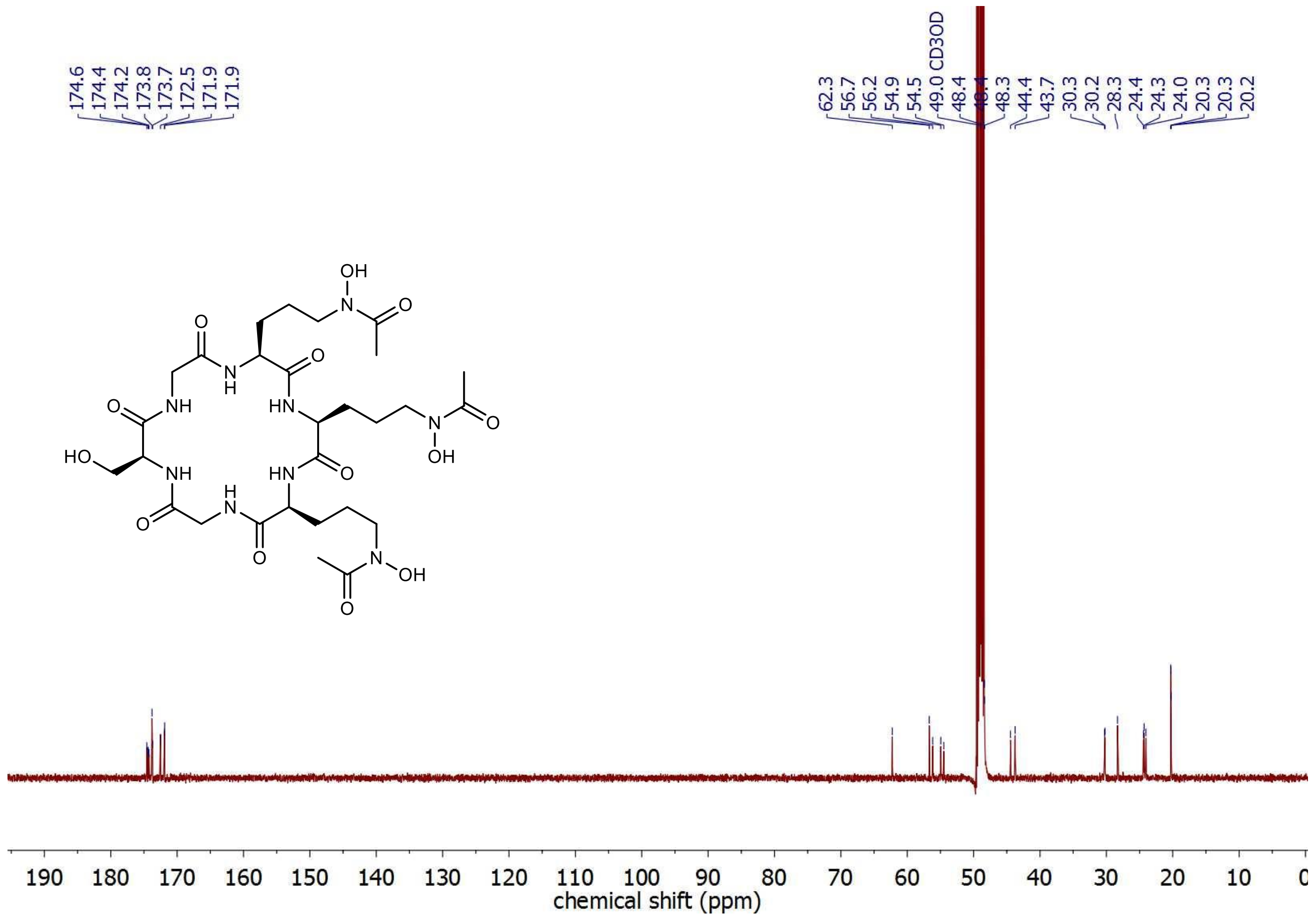


**Fig. S3: $^{13}C$ NMR spectrum of desferri-ferricrocin in $CD_3OD$.**

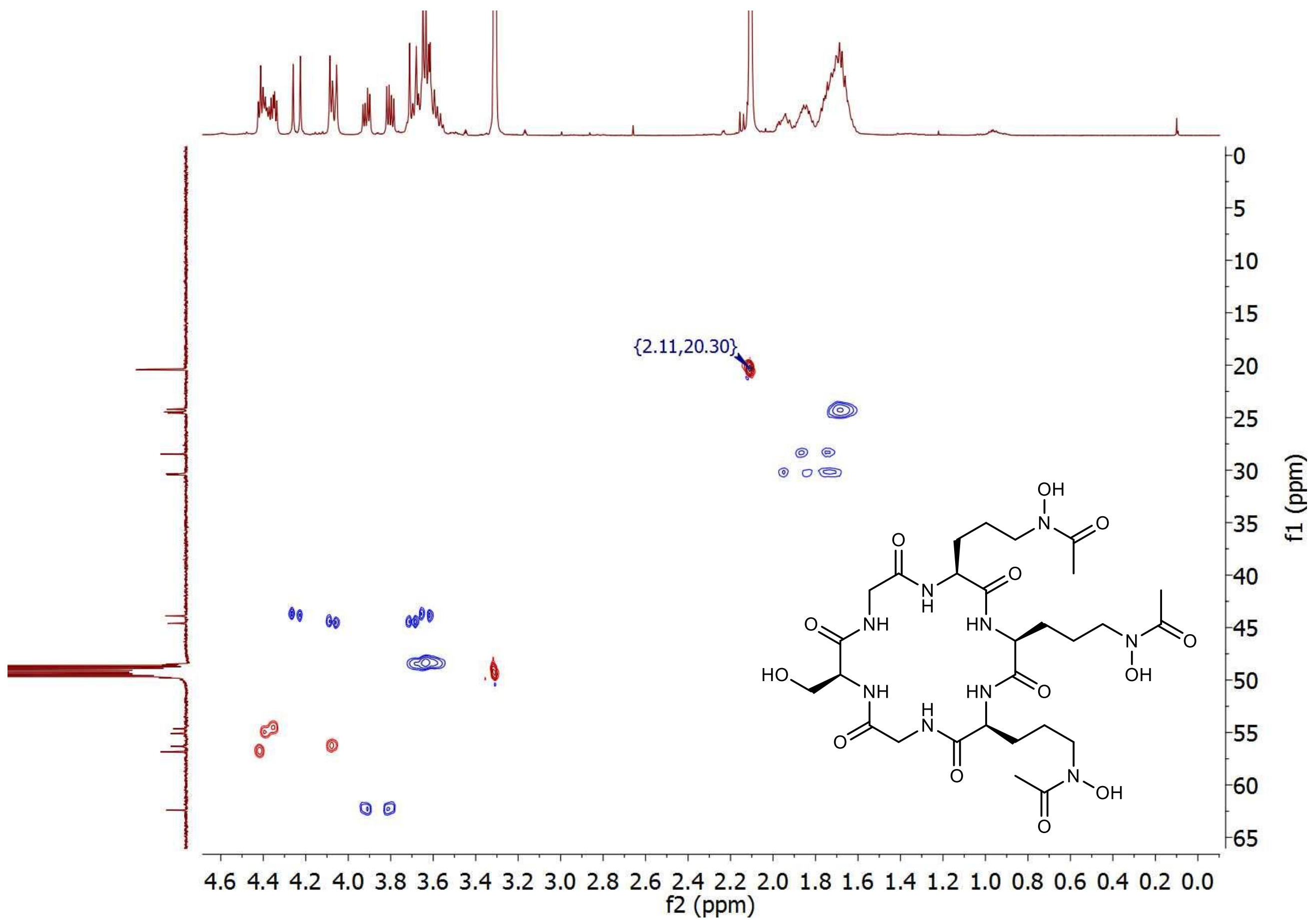


**Fig. S4: HSQC NMR spectrum of desferri-ferricrocin in $CD_3OD$.**

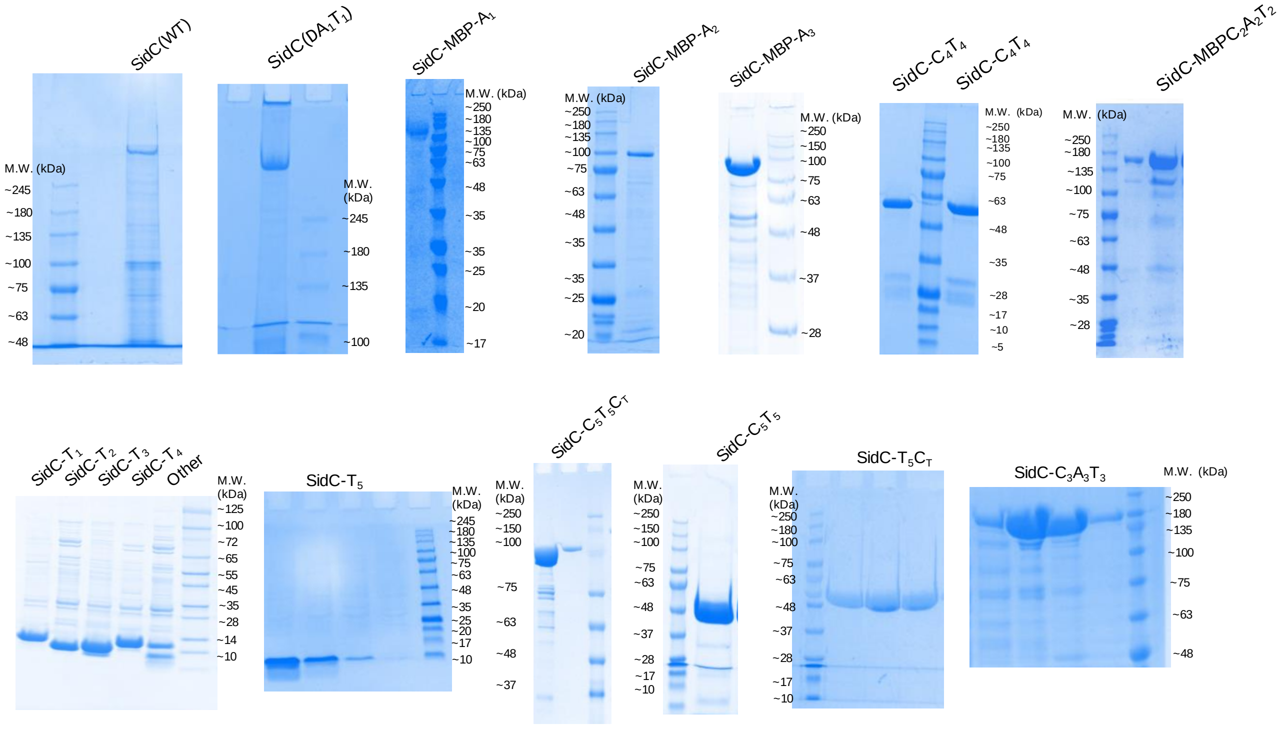

**Figure S5: SDS-PAGE analysis of recombinant SidC constructs used in this study.**

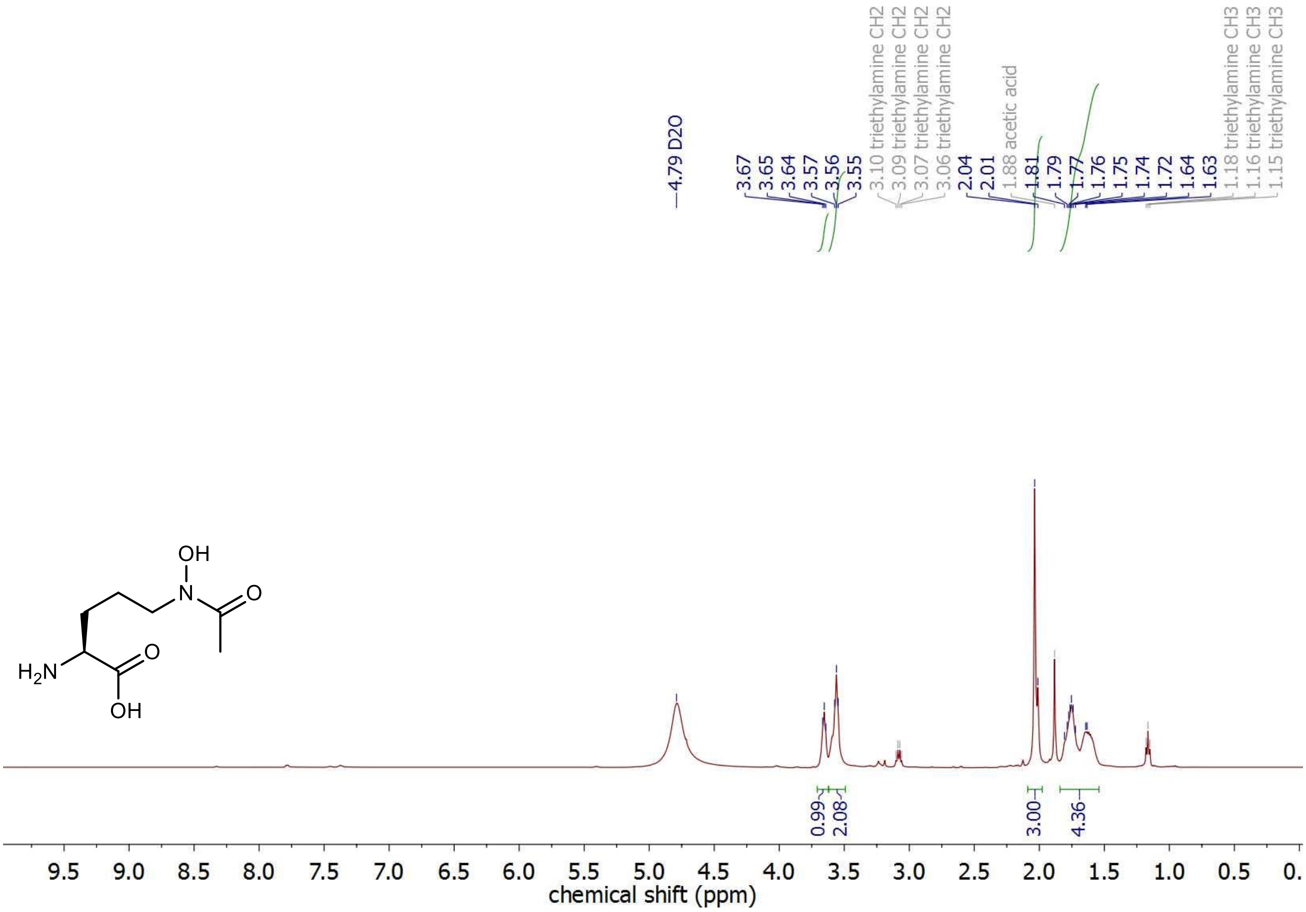


**Fig. S6. $^1$H-NMR spectrum of L-haOrn in $D_2O$**

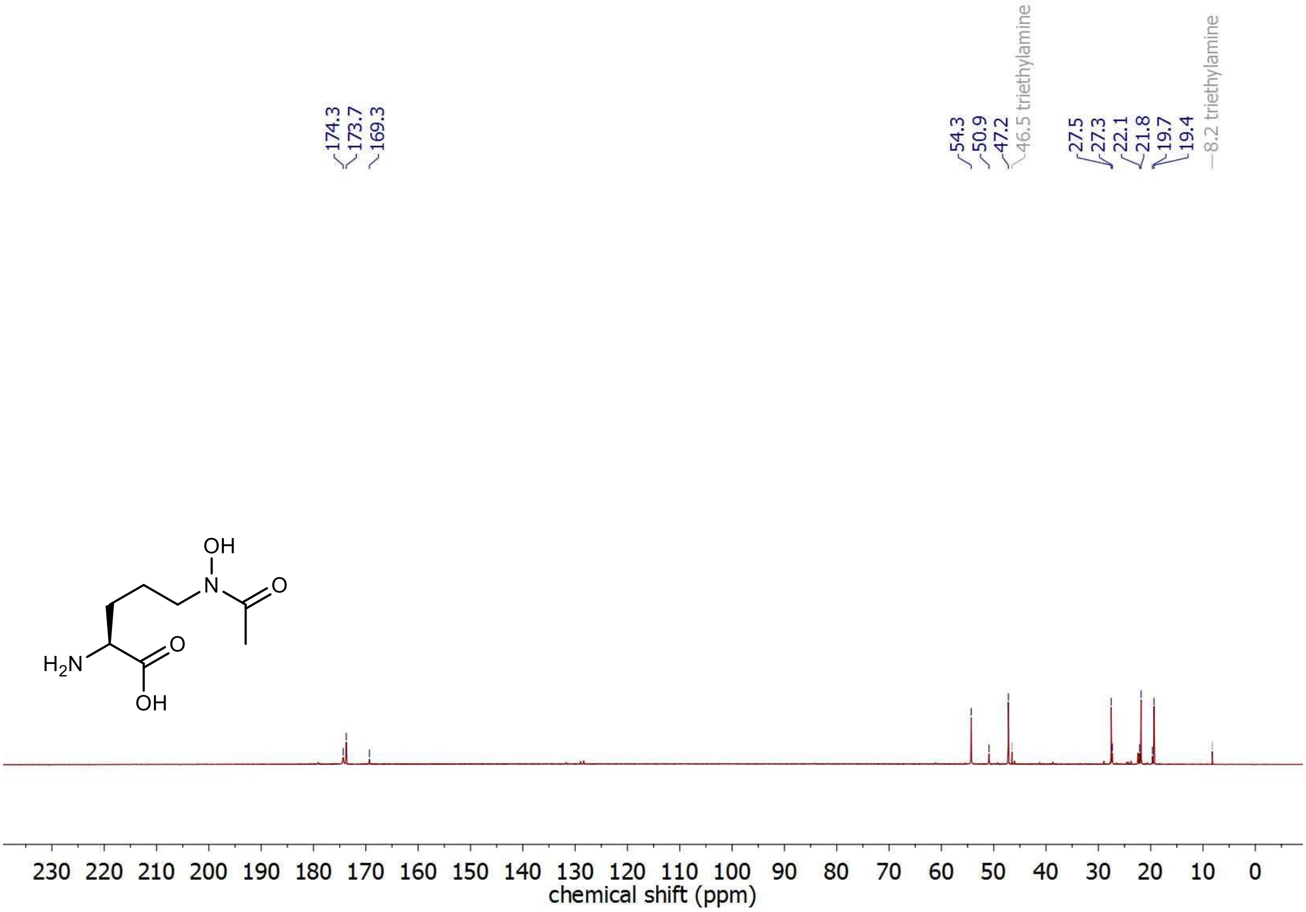


**Fig. S7. $^{13}$C-NMR spectrum of L-haOrn in $D_2O$.**

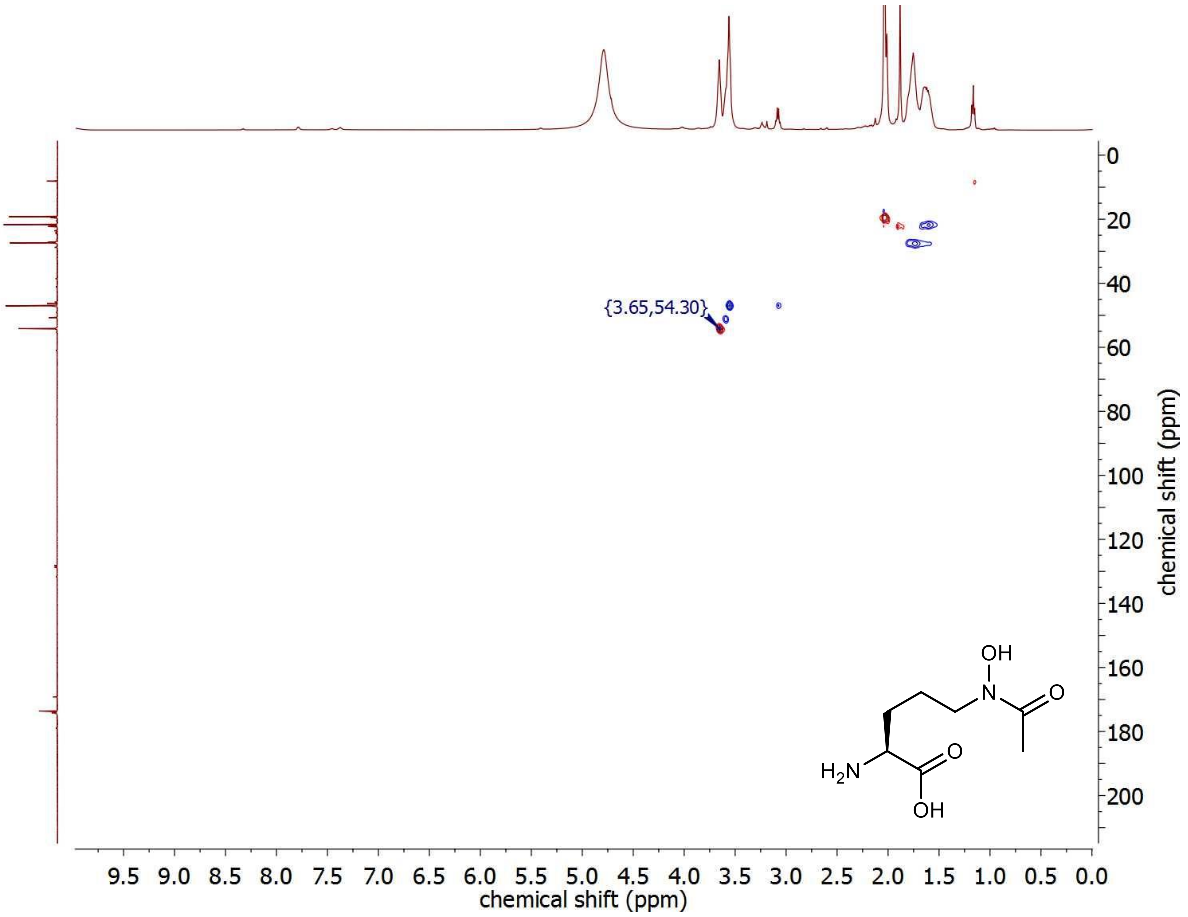


**Fig. S8. HSQC-NMR spectrum of L-haOrn in $D_2O$.**

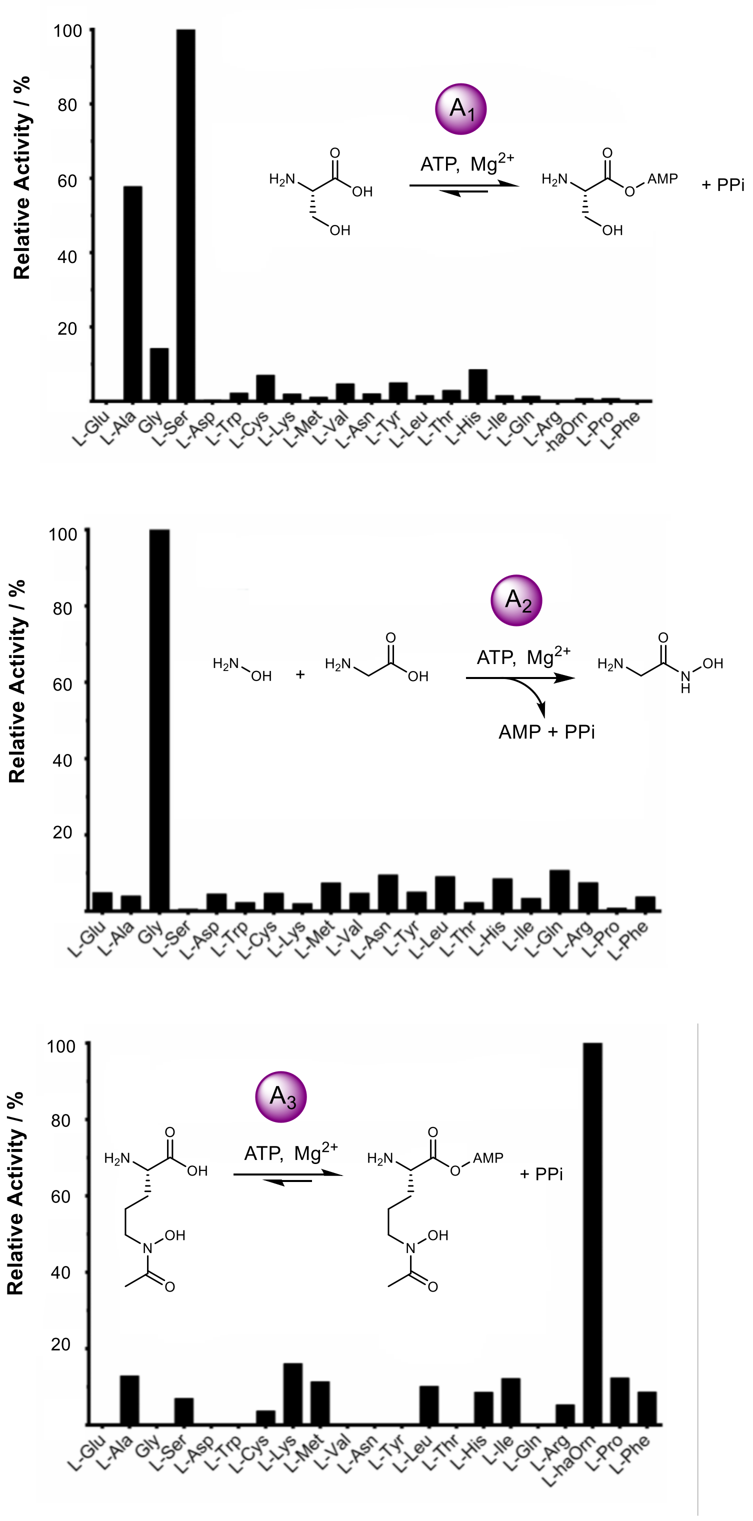


**Figure S9: Determination of SidC adenylation domain specificity.**
Analysis of SidC adenylation domain activity *in vitro* using ATP-[$^{32}$P]PPi exchange assay ($A_1$ and $A_3$ domains) and hydroxylamine release assay ($A_2$ domain) to determine substrate specificity. All values are displayed as relative activity normalised to: L-Ser (*top*, $A_1$ domain assay); Gly (*middle*, $A_2$ domain assay); L-haOrn (*bottom)*, $A_3$ domain assay. The constructs and conditions used for each assay are shown with each plot.

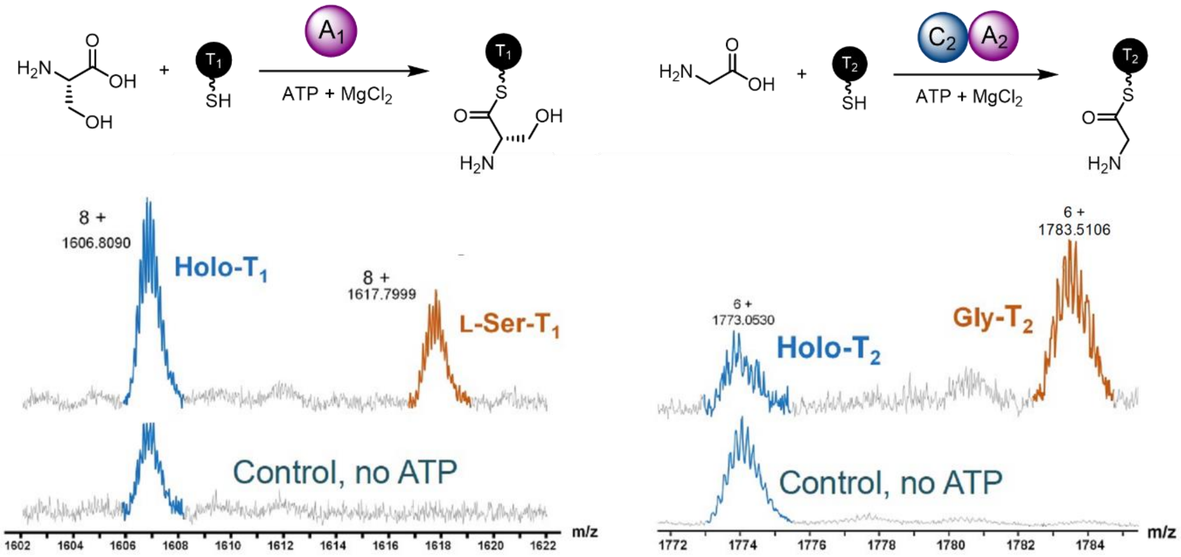


**Figure S10. Monitoring thiolation step of SidC $A_1$ and $A_2$ domains.**
Intact protein mass spectra of the SidC $T_1$ (left) and $T_2$ (right) domains following incubation with their cognate A domain, ATP, $Mg^{2+}$ and the requisite amino acid unit. The $8^+$ charge state of the $T_1$ domain is shown, with *holo-* and L-Ser loaded species highlighted in blue and orange, respectively. The $6^+$ charge state of the $T_2$ domain is shown, with *holo-* and Gly loaded species highlighted in blue and orange, respectively.

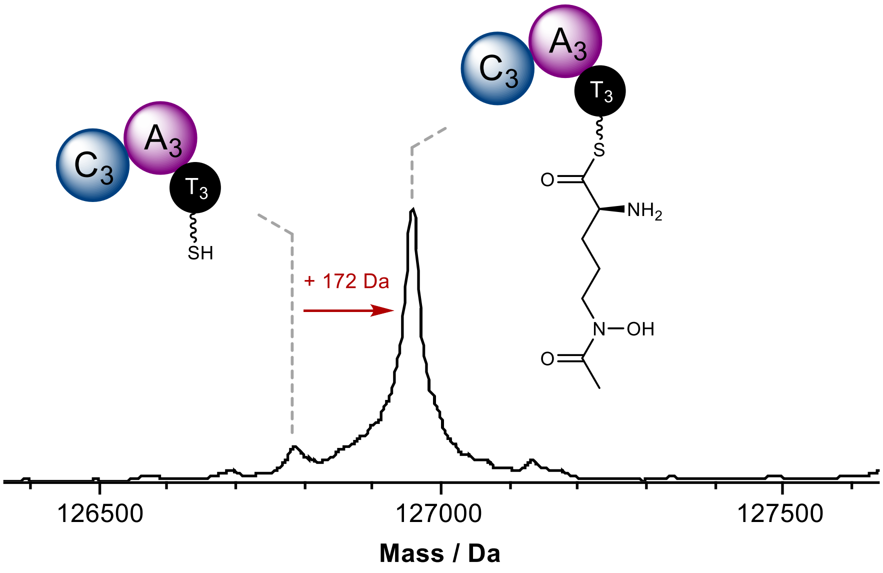


**Figure S11. SidC $A_3$ domain loads L-haOrn to the $T_3$ domain.**
Deconvoluted intact protein mass spectra of *holo*-SidC $C_3A_3T_3$ following incubation with L-haOrn, ATP and $Mg^{2+}$, showing loading of a single L-haOrn unit onto the $T_3$ domain, demonstrated by a +172 Da mass shift to the protein. Exact measured and observed masses are detailed in **Table S2**.

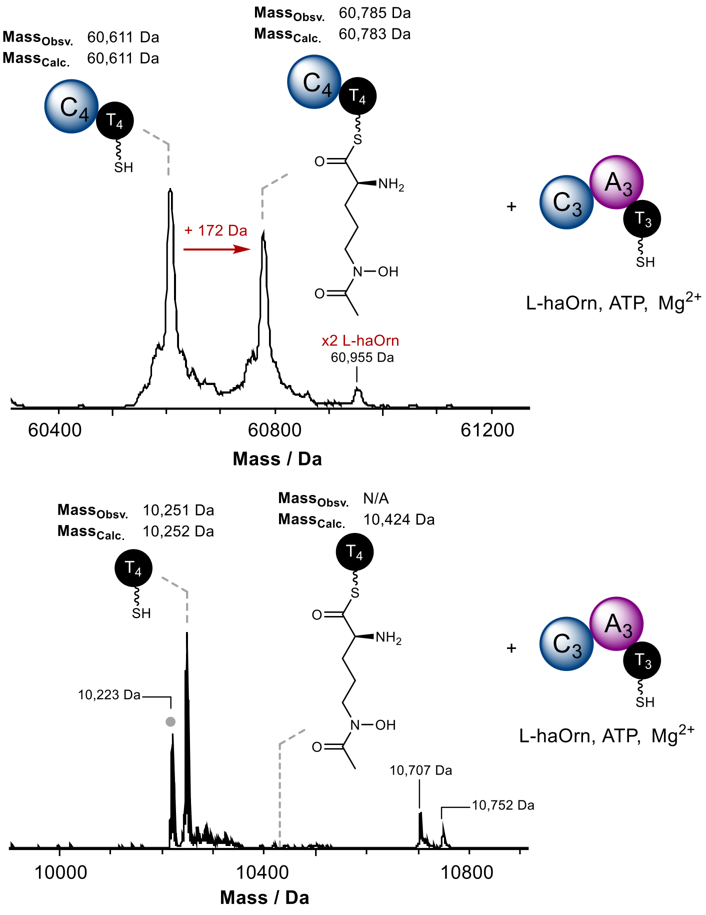


**Figure S12. Loading of L-haOrn by SidC $A_3$ domain to the $T_4$ domain requires the upstream C domain.** Deconvoluted intact protein mass spectra of *holo*-SidC $C_4T_4$ (*top*) and *holo*-SidC $T_4$ (*bottom*) following incubation with *holo*-SidC $C_3A_3T_3$, L-haOrn, ATP and $Mg^{2+}$. Loading of L-haOrn is only observed when the N-terminal C domain of each construct is present. Mass shifts corresponding to biosynthetic steps are highlighted with red arrows, and proposed intermediates are displayed. The grey dot indicates a species -28 Da less than the expected mass of SidC T4. The peak at 60,955 Da suggests trace amount of 2 x L-haOrn forming from condensation. Exact measured and observed masses are detailed in the spectra and **Table S2**.

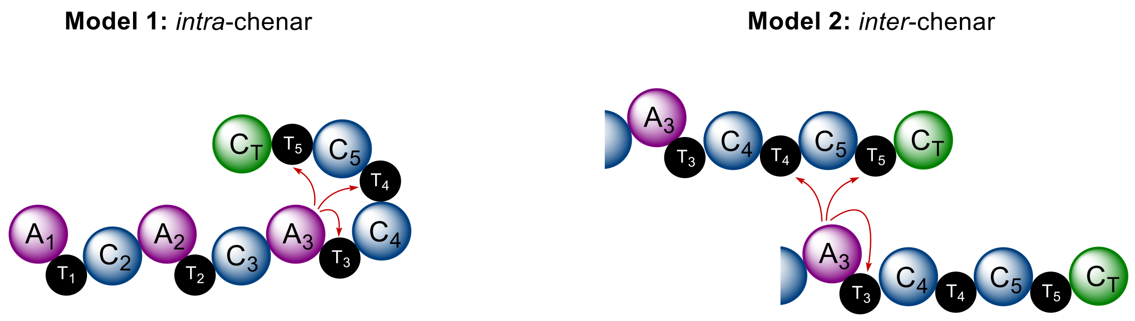


**Figure S13: Diagrams illustrating two architectural models for L-haOrn loading mediated by the $A_3$ domain.** In model 1, intra-chenar L-haOrn loading is facilitated by a 3-dimensional configuration that enables proximity of the $A_3$ domain to the $T_4$ and $T_5$ domains. In model 2, the juxtaposition of two SidC proteins allows the $A_3$ domain to load L-haOrn onto the $T_4$ and $T_5$ domain in an inter-chenar manner, whilst loading the $T_3$ domain conventionally.

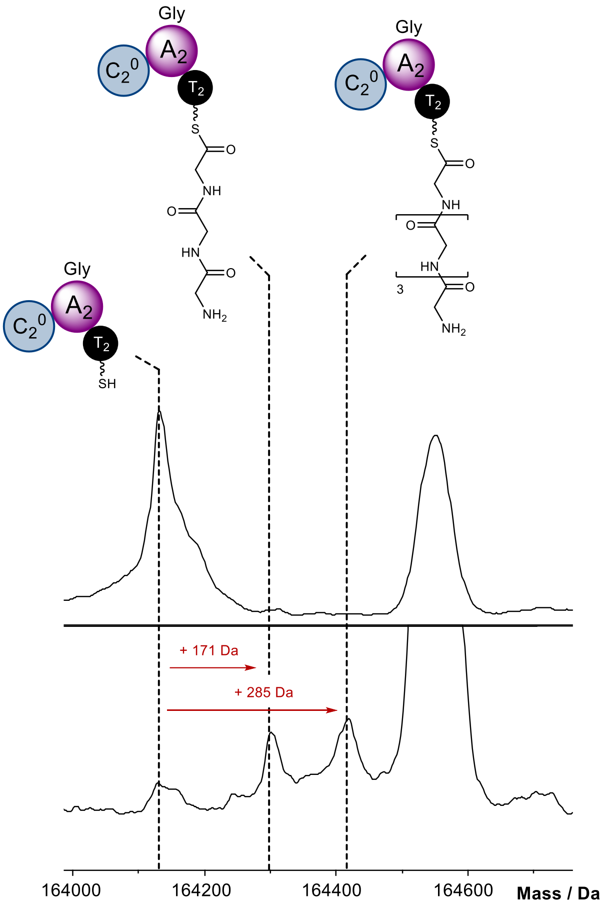


**Figure S14: SidC $C_2{}^0A_2T_2$ retains ability to produce $Gly_3$ and $Gly_5$ thioester intermediates.**
Deconvoluted intact protein mass spectra of *holo*-SidC $C_2{}^0A_2T_2$ (*top*) and *holo*-SidC $C_2{}^0A_2T_2$ (*bottom*) following incubation with Gly, ATP and $Mg^{2+}$. The spectrum shows a set of peaks corresponding to $Gly_3$ and $Gly_5$ intermediates in an identical fashion to **Fig. 3b, spectrum iii**. Note – the SidC C20A2T2 mutant gave reduced yields of protein, and therefore lower signal intensity in the MS. Mass shifts corresponding to biosynthetic steps are highlighted with red arrows, and proposed intermediates are displayed. Exact measured and observed masses are detailed in **Table S3**.

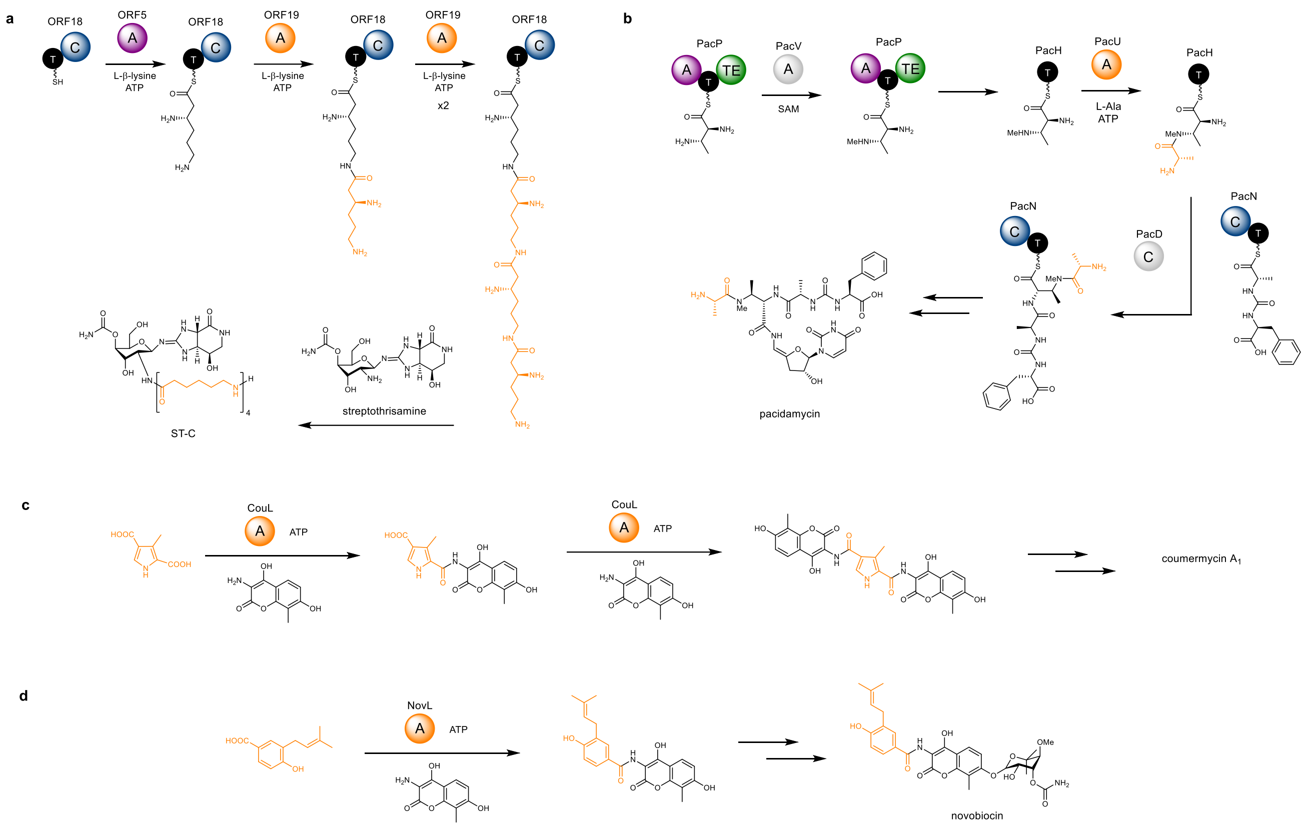


**Figure S15: Examples of other amide bond-forming adenylation domains in natural product biosynthesis.** Partial biosynthetic schemes highlighting key amide bond-forming steps conducted by adenylation domains for **a**). ST-C, **b**). pacidamycin, **c**). coumermycin $A_1$ and **d**). novobiocin. In all cases, the amide bond-forming adenylation domain and extender unit(s) are highlighted in orange.

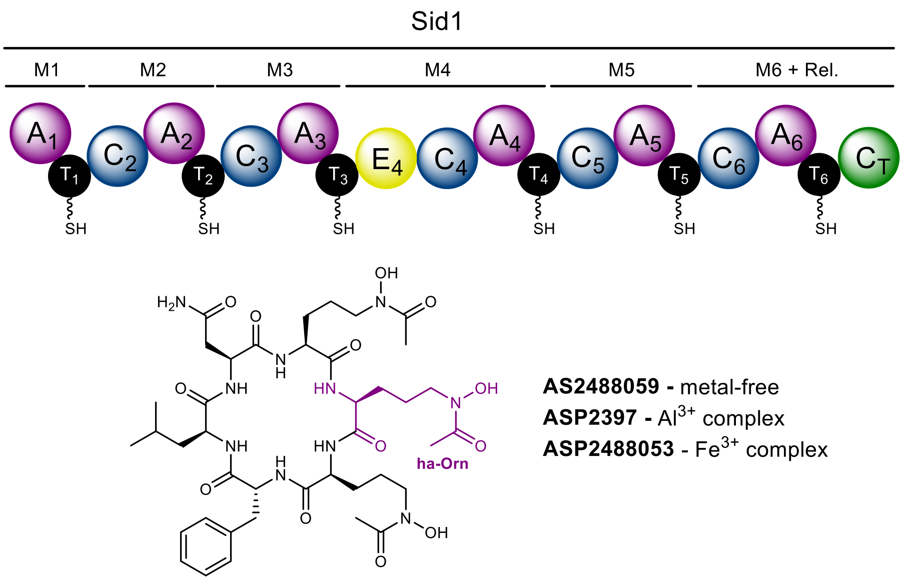


**Figure S16: Domain organization of the Sid1 NRPS responsible for biosynthesis of AS2488059.**

#### 2. Materials and Methods

##### 2.1 Molecular Cloning and Site Directed Mutagenesis

###### *2.1.1 Yeast Expression Constructs*

The SidC (AY223812.1), SidA (AN5823), and SidL (AN10080) gene exon fragments were cloned from the cDNA library prepared from the mRNA extract of *A. nidulans* FSGC A1145 strain[1] cultured on Czapek-Dox (CD) agar. The corresponding yeast expression plasmids were assembled through yeast homologous recombination using a Frozen-EZ Yeast Transformation II Kit (Zymo research). Gene fragments were integrated into a 2µ-based yeast expression vector with auxotrophic markers and ADH2 promoter and terminator regions. All proteins were cloned in-frame with an N-terminal $pHis_8$ tag to facilitate purification.

###### *2.1.2 E. coli Expression Constructs*

For bacterial expression, target regions of SidC were subcloned into either $pHis_6$-MBP-pET28a or $pHis_6$-pET28a vectors. All proteins were cloned in-frame with an N-terminal TEV-cleavable tag (either MBP or $pHis_6$), allowing removal post-purification.

##### 2.2 Protein Overproduction and Purification

###### *2.2.1 Yeast Expression Constructs*

The full-length proteins were expressed in *S. cerevisiae* JHY686[2] strain cultured in YPD medium. Briefly, single colonies of yeast cells harbouring expression plasmids was inoculated into SDCt uracil drop-out culture and left grown at 28 °C for 2 days. The seed culture was then inoculated into YPD culture (20 ml to 1000 mL) and left growing at 28 °C for another 2 days. Cells were harvested by centrifugation and washed once with cell lysis buffer (50 mM $K_2HPO_4$ (pH 7.5), 10 mM imidazole, 300 mM NaCl, 5% glycerol). Cells were flash frozen in liquid nitrogen and lysed by using a stainless-steel Waring blender. The cell lysate was cleared by centrifugation at 26,000 g for 60 min at 4 °C and the supernatant was filtered through a 0.22 µm filter (Millipore). The filtrate was incubated with $Ni^{2+}$-NTA resin for 30 min at 4 °C and then the slurry was loaded onto a gravity column. The resin was washed and eluted with increasing concentrations of imidazole in cell lysis buffer. The fractions were examined by SDS-PAGE gels. Pure fractions were concentrated to ~20 mg/mL by Amicon concentrators (Millipore), supplemented with 10 % glycerol and stored at -80 °C. Protein concentrations were determined by Bradford assay. Typically, 2 L cell culture could yield 1 - 10 mg of protein depending on the nature of the protein construct.

###### *2.2.2 E. coli Expression Constructs*

A single colony of *E. coli* BL21 (DE3) that had been transformed with the appropriate expression vector was picked and used to inoculate LB medium (5 or 10 mL) containing kanamycin (50 µg/mL). The resulting culture was incubated overnight at 37 °C and 180 rpm then used to inoculate LB medium (0.5 or 1 L) containing kanamycin (50 µg/mL). The resulting culture was incubated at 37˚C and 180 rpm until the optical density of the culture at 595 nm reached 0.6, then IPTG (1 mM) was added and growth was continued overnight at 15 °C and 180 rpm. The cells were harvested by centrifugation (4,000 x g, 15 min, 4 ˚C) and re-suspended in buffer (20 mM Tris-HCl, 100 mM NaCl, 20 mM Imidazole, pH 7.4) at 10 mL/L of growth medium then lysed using a Constant Systems cell disruptor. The lysate was centrifuged (37,000 x g, 30 min, 4°C) and the resulting supernatant was loaded onto a HiTrap FF Chelating Column (GE Healthcare), which had been pre-loaded with 100 mM $NiSO_4$ and equilibrated in re-suspension buffer (20 mM Tris-HCl, 100 mM NaCl, 20 mM Imidazole, pH 7.4). Proteins were eluted in a stepwise manner using re-suspension buffer containing increasing concentrations of imidazole – 50 mM (5 mL), 100 mM (3 mL), 200 mM (3 mL) and 300 mM (3 mL). The presence of the protein of interest in fractions was confirmed by SDS-PAGE, and an additional gel filtration step (Superdex 75/200, GE Healthcare) was used to further purify proteins where necessary. Fractions containing the protein of interest were pooled and concentrated to 250 - 400 µM using a Viva-Spin centrifugal concentrator (Sartorius) at an appropriate MWCO. Samples were snap-frozen in liquid $N_2$ and stored at -80 °C.

##### 2.3 Siderophore Isolation / Preparation

Desferriferricrocin and ferricrocin were obtained through coexpression of *sidC*, *sidA*, and *sidL* genes in *S. cerevisiae* BJ5464-npgA strain.[3] Briefly, competent yeast cells were transformed with plasmids XW55-SidC,

XW06-SidL and XW02-SidA and the colonies harbouring these three plasmids were selected using minimal medium dropping out uracil, tryptophan, and leucine. The colony was inoculated into the corresponding liquid minimal medium, and the cell culture was grown at 28 °C for 2 days. To induce production, the starting culture was inoculated to YPD medium and left grown at 28 °C for 3 days. The cell pellet was harvested through centrifugation and the produced siderophore compound was extracted using acetone. The organic extract was dried using rotavap and the residue was dissolved in methanol and subject to the LC-MS analysis on a Shimadzu 2020 LC-MS (Phenomenex Kinetex, 1.7 µm, 2.0 x 100 mm, C18 column) using positive and negative mode electrospray ionization with a linear gradient of 5 - 95% MeCN - $H_2O$ supplemented with 0.1% (v/v) formic acid in 15 min followed by 95 % MeCN for 3 min with a flow rate of 0.3 mL/min. To convert desferriferricrocin into ferricrocin, $FeCl_3$ (final concentration at 1 mM) was added into the organic extract.

To purify the fermentation product for structural analysis, similar extraction procedure was performed on 4 L cell culture pellet. The organic extract was dried and dissolved in $H_2O$ and fractionated with Amberlite XAD-16 (Sigma-Aldrich) resin. The desferriferricrocin and ferricrocin were eluted from a gradient from 20 % MeOH to 70 % MeOH. The eluent was combined and purified by semipreparative HPLC using a reverse-phase column (Phenomenex Kinetics, C18, 5 µm, 100 Å, 250 x 4.6 mm). The identity of desferri-ferricrocin was confirmed by HR-MS and NMR analysis. The NMR spectra data are consistent with the literature data.[4]

**$^1$H-NMR** (500 MHz, $CD_3OD$): δ 8.36 (s,), 4.46 – 4.30 (m, overlap, 3H), 4.24 (d, *J* = 17.1 Hz, 1H, Gly $C_\alpha H_2$), 4.08, (m, overlap, 1H, Ser $C_\alpha$H), 4.07 (d, *J* = 16.0 Hz, 1H, Gly $C_\alpha H_2$), 3.86 (ddd, *J* = 56.5, 11.1, 5.4 Hz, 2H Ser $C_\beta H_2$), 3.69 (d, *J* = 15.7 Hz, 1H, Gly $C_\alpha H_2$), 3.62 (d, *J* = 17.0 Hz, 1H, Gly $C_\alpha H_2$), 3.76-3.53 (m, overlap, 6H), 2.11 (s, 3H, hydroxamic $CH_3$), 2.105 (s, 3H, hydroxamic $CH_3$), 2.102 (s, 3H, hydroxamic $CH_3$), 2.00-1.90 (m, 1H, Orn $C_\beta H_2$), 1.90-1.85 (m, overlap, 1H, Orn $C_\beta H_2$), 1.85-1.80 (m, overlap, 1H, Orn $C_\beta H_2$),1.80-1.75 (m, overlap, 1H, Orn $C_\beta H_2$), 1.78-1.73 (m, overlap, 1H, Orn $C_\beta H_2$), 1.75-1.70 (m, overlap, 1H, Orn $C_\beta H_2$), 1.74-1.60 (m, overlap, 6H, Orn $C_\beta H_2$).

**$^{13}$C-NMR** (125 MHz, $CD_3OD$): δ 174.6 C, 174.4, 174.2, 173.8 (overlap), 173.7, 172.5, 171.90, 171.88, 62.3 (Ser $C_\beta$), 56.7(Orn$_\alpha$), 56.2 (Ser $C_\alpha$), 54.9 (Orn $C_\alpha$), 54.5 (Orn $C_\alpha$), 48.41 (Orn $C_\delta$), 48.39 (Orn $C_\delta$), 48.3 (Orn $C_\delta$), 44.4 (Gly $C_\alpha$), 43.7(Gly $C_\alpha$), 30.3 (Orn $C_\beta$), 30.2 (Orn $C_\beta$), 28.3 (Orn $C_\beta$), 24.4 (Orn $C_\gamma$), 24.3(Orn $C_\gamma$), 24.0(Orn $C_\gamma$), 20.31 (hydroxamic $CH_3$), 20.29 (hydroxamic $CH_3$), 20.22(hydroxamic $CH_3$). HRMS: calc. for $[M+H]^+$ $C_{28}H_{48}N_9O_{13}^+$, 718.3367, found 718.3365.

#### 2.4 Synthesis of L-haOrn Amino Acid Substrate

The amino acid $N^5$-acetyl-$N^5$-hydroxy-L-ornithine (L-haOrn) is synthesized from $N^2$-Cbz-$N^5$-Boc-L-ornithine according to the literature.[5,6]

**$^1$H-NMR** (500 MHz, $CD_3OD$): δ 3.65 (t, *J* = 6.0 Hz, 1H, $C_\alpha$H), 3.56 (t, *J* = 6.7 Hz, 2H, $C_\delta H_2$, *cis-trans* isomers not resolved), 2.04 (s, 3H, acetyl $CH_3$, with a small shoulder at 2.01 due to *cis-trans* isomerization), 1.83-1.56 (m, overlap,.4H, $C_\beta H_2$, $C_\gamma H_2$).

**$^{13}$C-NMR** (125 MHz, $CD_3OD$): δ 174.3 (carboxylate), 173.7 (amide carbonyl), 169.3 (amide carbonyl, minor isomer), 54.3 ($C_\alpha$), 50.9 ($C_\delta$), minor isomer), 47.2 ($C_\delta$ , major isomer), 27.5 ($C_\beta$), 27.3 ($C_\beta$, minor isomer) 22.1 ($C_\gamma$), minor isomer), 19.7 (acetyl $CH_3$, minor isomer), 19.4 (acetyl $CH_3$). HRMS: calc. for $[M+H]^+$ $C_7H_{15}N_2O_4^+$, 191.1027; found 191.1097.

#### 2.5 Biochemical Characterisation of SidC *in vitro*

Purified SidC and associated variants / mutants were converted to their *holo*- form by incubation in 20 mM Tris HCl, 100 mM NaCl, 2 µM of NpgA, 0.1 mM CoA and 10 mM $MgCl_2$ in a total volume of 50 µL for 1 hrs at 25 °C. Reactions were initiated by addition of ATP (5 mM) and either all or various combinations of the following: L-haOrn (1 mM) / L-Ser (1 mM) / Gly (1 mM) in a final volume of 50 µL, and the reaction was allowed to proceed at 25 °C. At different time points, the reaction was quenched by mixing with equal volume of methanol. The reaction products were analysed on an UHPLC-MS on a Shimadzu 2020 EVLC–MS (Phenomenex kinetex, 1.7 µm, 2.0 x 100 mm, $C_{18}$ column) using positive and negative mode electrospray S5 ionization with a linear gradient of 5 − 95% MeCN−$H_2O$ supplemented with 0.1 % (v/v) formic acid in 15 min followed by 95 % MeCN for 5 min with a flow rate of 0.3 mL/min.

#### 2.6 Biochemical Assays to Determine Adenylation Domain Specificity

##### *2.6.1 ATP-PPi Exchange Assays*

The amino acid substrate specificity profiles for the SidC $A_1$ and SidC $C_3A_3$ constructs were conducted using ATP-PPi exchange assays. Assays performed in 100 µL of reaction buffer (50 mM Tris-HCl pH 8, 2 mM $MgCl_2$) containing 1 mM TCEP, 5 mM ATP, 1 mM tetrasodium pyrophosphate ($Na_4PPi$), 5 mM substrate, and 5 µM enzyme. Before the addition of enzyme, $Na_4[^{32}P]$-PPi was added to a final intensity of ~2.5 × 106 cpm/mL. Reactions were allowed to proceed for 2 h at 25 °C and then quenched by the addition of 500 µL of charcoal (3.6% w/v activated charcoal, 150 mM $Na_4PPi$, 5 % $HClO_4$). Samples were centrifuged, and supernatant was discarded. To remove residual free $[^{32}P]$PPi, the pellet was washed twice with wash solution (0.1 M $Na_4PPi$, 5% HClO4). The pellet was resuspended in 500 µL of water and added to scintillation fluid at a final volume of 5 mL. Radioactivity was measured using a Beckman LS 6500 scintillation counter.

##### *2.6.2 Hydroxylamine Release Assays*

The hydroxylamine-trapping assay for detecting adenylation activity was conducted for the SidC $C_2A_2$ construct, and performed according to a reported protocol.[7] Briefly, the reaction was initiated by mixing 150 µL of substrate mixture [50 mM Tris, (pH 8.0), 30 mM $MgCl_2$, 300 mM hydroxylamine (pH 8.0), 10 mM carboxylic acid substrate] with equal volume of enzyme mixture [100 mM Tris (pH 8.0), 20 mM ATP, 20 µM enzyme]. For some hydrophobic substrates, 2 - 5% (v/v) DMSO was included to facilitate dissolving the substrate. The reaction mixture was then incubated at 30 o C for 16 hrs. The reaction was stopped by mixing with 300 µL of stopping solution [10% (w/v) $FeCl_3{\bullet}6H_2O$ and S5 3.3% TCA dissolved in 0.7 M HCl]. The precipitated enzymes were removed by centrifugation at 17,000 xg for 5 min, and 200 µL of the supernatant were transferred to a 96-well plate and the absorbance of the ferric-hydroxamate complex at 540 nm was measured by using a Tecan M200 plate reader.

#### 2.7 Biochemical Characterisation of *intra*-Molecular L-haOrn Loading by SidC $A_3$ Domain

Purified SidC $C_3A_3T_3$ / $C_3A_3T_3C_4T_4$ and $C_3A_3T_3{}^0C_4T_4$ proteins were converted to their *holo*- form by incubation in 20 mM Tris, 100 mM NaCl, 2 µM Sfp PPtase, 1 mM CoA and 10 mM $MgCl_2$ in a total volume of 50 µL for 1 hr at 25 °C. Loading of L-haOrn was initiated by addition of ATP (5 mM) and L-haOrn (1 mM) in a final volume of 50 µL, and the loading reaction was allowed to proceed for 1 hr at 25 °C before intact protein analysis by UHPLC-ESI-Q-TOF-MS (see **Section 2.11**).

#### 2.8 Biochemical Characterisation of *inter*-Molecular L-haOrn Loading by SidC $A_3$ Domain

Purified SidC $C_3A_3T_3$ / $C_4T_4$ / $T_4$ / $C_5T_5C_T$ and $T_5C_T$ proteins were converted to their *holo*- form by incubation in 20 mM Tris, 100 mM NaCl, 2 µM Sfp PPtase, 1 mM CoA and 10 mM $MgCl_2$ in a total volume of 50 µL for 1 hr at 25 °C. Loading of L-haOrn was initiated by addition of ATP (5 mM) and L-haOrn (1 mM) to a solution of *holo*-$C_3A_3T_3$ (100 µM) and one of *holo*-$C_4T_4$ / $T_4$ / $C_5T_5C_T$ and $T_5C_T$ (100 µM). The loading reaction was allowed to proceed for 1 hr at 25 °C before intact protein analysis by UHPLC-ESI-Q-TOF-MS (see **Section 2.11**).

#### 2.9 Biochemical Characterisation of *intra*-Molecular Gly Loading by SidC $C_2A_2T_2$

Purified SidC $C_2A_2T_2$ was converted to its *holo*- form by incubation in 20 mM Tris, 100 mM NaCl, 2 µM Sfp PPtase, 1 mM CoA and 10 mM $MgCl_2$ in a total volume of 50 µL for 1 hr at 25 °C. To a solution of SidC $C_2A_2T_2$ (100 µM), loading of Gly was initiated by addition of ATP (5 mM, or limited to a 2:1, 4:1 ratio with protein) and Gly (1 mM, or limited to a 2:1, 4:1 ratio with protein) in a total volume of 50 µL at 25 °C. Loading reactions were allowed to proceed for various time intervals before intact protein analysis by UHPLC-ESI-Q-TOF-MS (see **Section 2.11**).

##### 2.10 Biochemical Characterisation of *inter*-Molecular Condensation Reaction Between SidC $C_2A_2T_2$-$Gly_3$ and SidC $C_3A_3T_3$-L-haOrn

Purified SidC $C_2A_2T_2$ and SidC $C_3A_3T_3$ proteins were converted to their *holo*- form by incubation in 20 mM Tris, 100 mM NaCl, 2 µM Sfp PPtase, 1 mM CoA and 10 mM $MgCl_2$ in a total volume of 50 µL for 1 hr at 25 °C. Reactions were initiated by addition of ATP (5 mM), L-haOrn (1 mM), Gly (1 mM) to a solution containing *holo*-$C_2A_2T_2$ (50 µM) and *holo*-$C_3A_3T_3$ (100 µM) in a total volume of 50 µL at 25 °C. Reactions were allowed to proceed for either 10 min or 60 min before intact protein analysis by UHPLC-ESI-Q-TOF-MS (see **Section 2.11**). A variation of this reaction was conducted which allowed $Gly_3$ and $Gly_5$ to be formed *in situ* on SidC $C_2A_2T_2$ (following procedure of **Section 2.9**) before its addition (at a final concentration of 50 µM) to a solution containing *holo*-$C_3A_3T_3$ (100 µM), ATP (5 mM), L-haOrn (1 mM).

##### 2.11 UHPLC-ESI-Q-TOF-MS Analysis of Intact Proteins

Biochemical assays were analysed on a Bruker MaXis II ESI-Q-TOF-MS connected to a Dionex 3000 RS UHPLC fitted with an ACE $C_4$-300 RP column (100 x 2.1 mm, 5 µm, 30 °C). The column was eluted with a linear gradient of 5 – 100% MeCN containing 0.1% formic acid over 30 min. The mass spectrometer was operated in positive ion mode with a scan range of 200 – 3000 *m/z*. Source conditions were: end plate offset at −500 V; capillary at −4500 V; nebulizer gas ($N_2$) at 1.8 bar; dry gas ($N_2$) at 9.0 L $min^{-1}$; dry temperature at 200 °C. Ion transfer conditions were: ion funnel RF at 400 Vpp; multiple RF at 200 Vpp; quadrupole low mass at 200 *m/z*; collision RF at 2000 Vpp; transfer time at 110.0 µs; pre-pulse storage time at 10.0 µs. Measured masses for all species are displayed in **Table S2** and **S3**.

#### 3. Sequences and Tables

Numbering of amino acids excludes the His-Tag regions (highlighted in grey), and correlates to numbering referenced in the main text.

#### ***sidC*** **NRPS (DNA)**

ATGGCTAGCCATCACCATCACCATCACCATCACACTGCAATGGGGAAACGTAAGCTGGCTGGCTCGTTCCGCAGATATGTCGCTACCGGGCTTTCTCGGGTCTCGGGCGCTTTTCGGAGTCGCAGT
GCCAGTAAGGAACGCCAGAGCCGTGGCAGCGGCAGCTTGGTCGAACGTGATACGCTGGCAAAAGACCAGGGCCAGGGGGTCTGTTTACAGCCAGCTCAAGTTGGAGCTGTCCCTGTCACTAAGGT
CCCGGCGCTTGATACGATCGAAGCATATGCCGGAGCATTTGATAAACCAATTGACGGGGCCGGTACGTGCCAGCGGACAAGTTTGGTCTGCAGTGCCAGCTCTGCCGTTGACGGTGACGTGATTG
ACGGCTTGGTTCAGGGCTATGCGCGGTTCATCGCTGGATTGACGGGTCTTGACGATATCGCGTTCTACTCCACTCGTCATGAACCGTTTGCATTGGAGAAGTCGACATCAATCATTATCGCTTCTGC
TGCTTCCAACGAGTTAGTCTGTCGTGAGGTGGACACGAACCACGAGAACGATGTGCAGTTCTATGCTGAGCTGGGACGGGTCGAGCCTGAGAATGGACGACGTGAATTCCAGCCGAATGCATTCAC
ACTCTTCATCGAGCCGGACAGCAATAGGAAGAAGAAGAATGTTCTTCACCTTAGCTTCGCCTATCCTCGACGGCTGATCCCAGATGCAGCGGTAGAACAGCTACTGCGCACGCTTTTACTGCACATC
TGCGAGTCTTCTCCACTAAAGTCGCCATCGACATCGCAGCCTGAATTGTCCATTCTGAATTTCCCGCCGTCTATGATACCCCCAACTGCGCAGGCGAACGGCGTCGAGAATTCGACGACCAACCCG
CATCTCTTGCATTCCGCATTCGAGAACTGGGCTCGCAAGAACCCGCATTTTATCGCATTAGATTTCATTCATTCCCTTTCATCCAAAACGCACAGATCAGAGCATAGCATCATCACTTATGCGGCCCTC
GATAGCGCTGCTACCAATCTAGCCTTCCATATTAGATCACTTCTACGCGACTCTCGAAAACATGGACAGATTATCCCTGTTCACATGCCCACTTCACCGGAGCTCTACATCTCATACCTCGCCGTCCT
GAAAGCTGGCCACGCTTTCTGTCCTATACCTCAAGACGTTCCTGCCCGACGAATACAGGAGATTCTTAGCGATATTGATGCTCCTATTGTCCTCGGGACCTCATCAAAACCGCCCATCAGTGCTGAAA
GCTCCCGGTCGACGTCTACATGGGTTAATGTCACCGAAGTTTCCAAATGGCGGCAGATGTGTGGCGAGCAACCAGCTGATTATTCTCGACCCTCTTTAGACCACATAACCATTGAGCAGAACCAGAC
TGCTTATCTCCTCTTCACCAGCGGTTCAACCGGTAAGCCGAAAGGTGTGCAAATCAGCCATCTCGCAGCATCATGCTCCATATCCTCCCACGCCACTGCAATTCCTCTCCCCGGCGAATCACCAGGC
AACTTCCGCTGGTTCCAGTTTGCTTCGCCATCCTTCGACCCTTCACTCATGGAGATTTTTGTCACCCTAAGCACTGGTGGCACGCTCTGCTCTGCTGATCGCAGACTTACCCTCGCCAATCTCGAAGC
CACCATCAATGAGAGCAGAGCGACAGTCATGATGGCTACGCCGAGTCTTGCCACGCTTCTCAGACCTGACCGGCTGGAGACCCTTGAGGCGCTGTGGTCCATGGGAGAAAAGCTCAACCGGACAG
TTATTGACAACTTCGCGCTGGATAATGTTATGAATGGAGACGCCGAGACCAGAAGACCGAGGACCTTGGTCAATGCATACGGTCCAACGGAAGGAGCAATCAACTGTACCTACGTCGCTCCCTTCAA
GCGTCACATGCGTGGTTCTATCATTGGACGGCCGCTACCAACCTGTGCGATGTTTATTCTATCCCCCGACTCCCAAGTCCCTGTCTTAGTCCCTACCGGTACAGTTGGAGAACTGGCAATTGGCGGC
CCCCAAGTCAGCAAAGGCTATCTCAATCTCCCCGAAGTAACAGCTAGAGTCTTCATTCGCAGCAAGGAATTTGGACCCCTCTACCGCACTGGCGACAAGGCCCGCATCGTCTGGGACGAAAGCGGG
CACCAGGTCATTGAGTACCTAGGCCGAATCCGCACGGATCAGGTCAAGATAAATGGACGCAGAGTTGAGCTGGGTGAAATTGAGTCTGTTGTCGCGGCTGTGGAGGGTGTTCGGGAGGCCGTTGC
GGTTGTGGTAAAGAGAGATAGCAAGTCAAACGGGGGCGAGCAGATTGTCGCTTGTCTTGTGGTAGATGCTGCGGATGGGGAGGGGCGTGAGAAAATTGCACAGCAAGCAAAAAAAAATGCTGCGC
AGCATCTTGCGAGCTTCATGTGCCCGACTACCTACACATTTTTTGATGTGCTGCCCAGGTCGAGTTCCGGAAAGGTCGATCGAAAAGCGTTGGCTGTCCAGCTGCAAGAGAAGCCTGATGTTCCAAT
CAAGAATGGCCTCAGTGAAGAAAGTGCAGAATGTTGGAGGCATTCAGATGAGGCCGCATCGTCAGTACAGCAGCTGGTGATTCGACTCGTTGCGGAAACAGGGGACGTGAGTGATTCCGCAATAAA
GCCCGGGACAGAGCTGTACTCGGTAGGCATCGACAGTCTAGGAGCAATGCGTTTTCTACAGAAGCTGAGAGACAATGGGGTCCATGGGCTGTCCGTCGGCGAAGTGCTGCAAACACATACATGCC
AGCGGCTTGTCTCTCTCGTCCAGAGCATGCTGACGAATCAAAATGGCGAGCCTAATGGGATACTTCAGAAGGGCAGTATAACAAACGACTTGCAGTTAAGACTGCAGTCATTCGGCCGGCGATATCG
CATGTTTTGCGCAGAATCCCTTGACGTGCCCGCGGACACGATACATGAAGTCCTTCCTACCACTGCTACTCAATCCGGGATGTTGACTAGTTTTCTGCGTAGCTCTGCGGAACAGTCCTATGAGAAA
CCAACATACATTTATCATACTGTCCTTCCACTTGAGCCCCGTACCAACATTGAGAAGCTCAAGAAGGCCTGGTTCGATGTTATCTCAAACTATGACTCTTTCCGGACGGTCTTCTGTATGGTCGACGAT
GAGCTGGCCCCGTTTGCACAGTGCATCTTGACGGCTGAGGGCGTCTCATCCAGGGACTGGAACGTTTACACAAGCCCTAACGAGTGCTCCGCAGAAAACGACATTATCGACCACGCTCTTCGGAGT
GCTGAAGAATCAATCACATTGAGGAGACCTCCCTGGAAGCTGTCTTTGGTGCAGTCATCGACGAAGTCCATCATGATCCTCAGCATGTTCCACGGAATCTTCGACGGCGGCTCTCTACAGCTCCTCT
TGCAAGATGTCTCGTCAGCGTACTCCGGAAACACACTACCGCAGCGAACCTCACTAACCCACATCGTCAAACACCACTTCCAGGCAGATCACACTTCCACATCCAAATTCTGGAGGGAATATCTACA
AGGTTACTCTCTCCTCCCCTTTCCATCTCTCACTCCGCATCGAGCGCCGGCCCAGAAAAGCACTGGCTGTGCTGAGGTCACCTCTCAACTCAGTTACGGAGCCCTCAAAAAACTCTCAAAAAGCATT
GGCTCTACACCACTCTCTGTGCTGCAGGCAGCATGGGGTGCGGTAGTTCTCAGCTACACAGCTACGCCGGATCAGGACGTAGTTTTGGGAAGTGTGATGTCTGGACGACTTGATCCCGATTCGAAA
GACTGCATTGGTCCGACCTTTACGATAACGCCTTCCAGAATTTCCGTCCAGCAATTGAAGGGTTCCGGATCCTTGACAAACCAATCAGTGGTCCACTATCTGTCGTCTTCAAATGCAAAAGCGCTCTC
ACACCTCCAGCCTCAGCTAGGATCTGTGACCGTAAATGGCAAGGTTCCCTATGACACAGTCCTCGCATACCAGGACTTTGACACTGATATGAAACCTTCTCAAACCTGGAGTTCTGTCCAACATCCCG
CTATGGCAAATGATTTCTCTGTGATGATCGAGGTGGTCCCCAATCCGGACTCGACGTTGACGCTGCGGGCTAGTTTCGACACGAAACTCGACTCTACCGGAGCTCAAATCATGCTGAAACAAATGGA
TGATATCGTCAGCTACATCCTCAACCACCCGGACTCGAGTTTCGAAGATGCTCCTCTTCAAGCCAGCTTAGCTCTGAAATCGAAAGCGAACCCCAGTCCAATTACAGCTCCAGAAGTCAGCGAAGGA
GCACTCCTACAATCACAGTTCGAAGACCATGCCCTCTCGCATCCAAATGACCCAGCTCTAGTATTTAAGCAGGACCTTGATAATGATGATCATCCAGGAAACATCACTTGGACATATGCCCAACTCAA
CGCTATGGCTGAAGCTCTTGCGGAACATCTGCTCCAGGTTTGTGGTGATCTGAGAGATGCTTCTGTTCCGATTTGCATCGAGAAGAGTCCACCCTTGTATGTTGCGATCCTAGGTATCCTCAAGGCA
GGAGGCGCCTGGTGTCCCATTGATACACTTTCCCCAGCACAGCGTCGCCATGATCTCATTGCGCGAACTAGTGCCGGTATCCTTCTGGTTTCTGGTCTCGATACGCCACAACCTCAGAATGCTGTTC
CAGCTGGAGTCAGAGTTATTGATGTTAGCAAGTTCATTCAGAATGTCTCGAGCGACAATACCCCACAGTCAAGCAGACACCGAGCAACCCCTAGAAATACAGCCTACCTAATCTGGACGAGTGGCAC
AACTGGCGCTCCCAAGGGCGTACCCATTACGCACTCAGCCGCCGTCTCTAGCATGCGCTCCCTCCAGACCGATATCCCGGGTAACGAAGACGGCAGTCCCATTCGCTGCCTCCAATTCTCGCAACC
AACCTTCGATGTCTCCATCCAAGACCTCTTCTACACTTGGGGCTTAGGAGGCGCGCTTATTTCTGGCACGAGGGAGATAATGCTCGAATCTTTCCCTAAACTTGCAAATATCACCAAGGCAACGCACG
CCCATCTCACGCCTGCCTTTGCGGCTGGTGTCGCGAGGAAATCATGCAAAACGCTTAAAGTCGTAACAATGATCGGCGAAAAATTGACACAGAGTGTTGCGGATGATTGGGGGACAGACATGAGGG
CGTTCAATACCTACGGCCCCGCGGAGGCGACGGTCGTCAGTACGATCCGCGAATTCGGCAATGAACATAGATCCGTCAAAAGTGCGAATATCGGGTGGCCAATGAGCAGTGTTTCGGTCTTCGTGA
TGTCCAAAGATAAGCGCGTGCTCATGAAGAATGCCATTGGAGAGCTTGCATTAGGTGGCCCTCAGCTTTCACCAGAATACTTGAATTTAAAGGACGTAACGGACACGAAGTATATCTGGAATGAGGA
CGCTGGGCAGCGGCTTTACTATACGGGTGACCTTGTCCGCATGCTATCTGATGGGAGTCTGGAGTACATCACGCGTGTTGATGACCTTGTGAAACTAGGTGGCATTAGGATTGAGTTGTCGGAGATA
AGTTTTGCGCTGAGAGGGTGTCATGAACTCGTCGAGAGCGTGGAGACTATGATACTTAGCAGGAAAGACAGGCCAGTCAGAGTTGTTGTTGCGTTCTTGTGCGCCCCGAAGGCTGCTGGTGATGCG
GATGAAGGTTTGTTGGTGCTTGACGACACTGGTAGAGATATTGCCCGTGCAGCAAGCCTTCAGGCGCGCAATGTCCTGCCAGAGAACATGATTCCCTCTGTGTATCTTATTGTCAAGAAGATACCAA
AAACGCCATCTGCAAAGGTCGATCGACGCGCTTTACAGGCTGCGTACGCAGCGGTTGATATTGACAAGTGGGAGAACAACGTTAACCCTGAGGGCCCCGGAGATGCAGATGAGGATGATGCTGCC
ACGGCCACTCAGATCATCGAGACAGTGGCAGCGCTGGTGCATGTTGAGAGCTCGACTATCACAAAGAGCAATAGATTACGCAGTCTAGGTGTTGACTCACTCTGTGCCACGCGTCTTGCATTTAGGC
TCAAGGAGGCCGGGTTTGGTCTTTCTGTTATGGATGTCCTGGCCTGCACCACTATCCAGGATCTTGTCAGATTAGCCCAGTCTACTTCATTCAGTTCATCTGCAAACTCAGCTAGCCCTCACAACGAC
AAGTTCGATATCGTCTCCTTCAACAAGATATGGCATACCCTCGTCGCAGCAGCAGCAAAGATCCCTGAGAAGGACGCGTTTACAACGATCCGCGCCACAGCTATCCAGGAGAGCTTGCTTACTGAGA
CAATGGGTACATACGACATGTACTGGAGCAATCATTTCTTCAGACTTGATAGGTCTGTCGACATTCCCCGACTACGACAAGCTTGGTATGCCGTGTGCCAGAAGACTGAGACGCTCCGGACGGGGTT
TATACCTGTCGCACAGACGGAGGCAAAGAATAAGAAACAGGCTAAGGATTCGGGATTTTCTATTTTGCAGGTTGTGTATAAGCTTCCGGCGGTGGATTGGGAGGCTCATACCTACAGGAAGGAAGAG
TGGAGTTGGGTGCTAAAGAACCGAGTGCGGGATATCATGACGGCGCACCAGAAGAACTACTTCTGCCATCCTCCTTGGGCAGTCACAATACTGGAAGAAGGCACCGAACGAGTCATGGTCTTGACA
CTCCACCACTCAATCCATGACGAGCCTTCCCTCAAGTTCTTGATGGACGATGTCCGCGCCGCATATACCTACAAACCGCCTCTTCGCACCCAGCTGACCCCCGCCCTCGCGCTTGTGCTCCCAACTC
CAAGCAAATACGCCGAAGCTGTTGATTTCTGGTCCTTGGAGCTCAAGCCGTACGCCGCCCTCGACGTCCCCGTCTGGCCCGATCTCACAGGTAAACGGGTCTCACCAGGCGCGGCGCCAGAATAC
AAACTGATATCCGAGGCGATGTCAATAACCTCGTCCTTCCCAGTTTTAGAGAAAGTCGCAGCAGATCTAGGGCTGAAATCTGTTGCGTCGATCATTAGGGCGGCATGGGCGTTCGTCAGCCTGTCTT
ATCTGGGCCTGTCGGGGACGGTGTTTGCGGAGACGTTATCAGACCGCGTCTTTGATCCGTCCTTGGAAAGCGCAGTCGGTCCTTTTATATCTGTTGTTCCGGTTCCGTTCCGAGTAGAAGGGGATA
CGACAGTGAGGAAGATCCTGGCAGAACAGCACCGGCTCTCCTTGCAATCGTGGAAGCATCGACATGTTCATGCGAGAGATATTCGGAAGGCCCTTAAGCGGCAGAGAGGAGAGCCGTTGTACCCT
GCAGTATTCAATTTCCACGCGCTTGACGAATCAAAGGACAGAAAAATTGCGAGCCTGCCTGGTCTCTGGCATGAGCTTGAAGACCAGATTGGCCTGCATGTTGAGCATCCCATGGCCATGAATGTTT
TCCAGTCGCCAAGCGGAAACATGACCCTGGAGGCATCAAGCGATAGTCGGATCTTCAGCCGTGAACATTTACGCCTTTTTGTTCGCCAGATCGATGCTCTCATCTCGGAAATGCTCTTATCGCCTGA
TGAATCTCTCAGCGGGCTCGTGAATCGGCTTCCGTCTGACCTACGATCCCTATCCAACAGGATTGTGAGCCATGAAGTCGCGAATTCAATTCACCAGGCCCCAACTTACTGGCTGGAGAAGTTCGCC
GACACTCACCCGCACTGGACAGCCGTCGAGGTGGCGTCCAGTATCAGTACAAACGGAATTGAAAAAGAAGCCATGAGTTATGGCTCTCTAAACAGCGCAGCAAACCGTGTTGCTGCGTACTTGGCT
TCATTTAGGTACAAGAATCGAGTCGTCGGTGTCTGTGCAGGACGGACATTAGCGTCGTACCCCATCATCATTGGCATATTCAAGTCCGGGAATACATACCTTCCTATTGATGAGAGTCTGCCAGCCG
ATCGAAAGGCGTTCTTGCTGGAGGATGCAAAGTGTCCGGTTGTCTTTACGGAACTGGGTCTCAGGAATTCCTTTGCCGGTGCTCCTGATACCTGTCGCGTGGAGTGTATTGACGACCCTGCCCTGC
AGAGATCATTGGACGAAATGCCATCGACAAACAAAGACTACAGCTCCCATCCAGATGATGTCTCATACCTGCTATTCACTTCCGGCTCGACCGGCAAACCCAAAGGCGTCATGGTCACTCGCGCGAA
CCTGTCGTCCTTCATCGAGTCCATATCTGAGTTTGCATGCCGCATTGCACCAGATACGCTTAAGCTGGGTGGCACAGGTCGCTATCTAGCGCAAGCAAACCGCGCCTTCGACCCGCACCTGCTGGA
GATGTTCTTCCCATGGCGCCATGGCATGGCCACTGTGACTGCCCCGCGGCCTATGATCCTTGATGATATCGGGACGACTCTATCCAAATGGAGCATTACACACGCCAGTTTCGTTCCCAGTCTCGTC
GACCAGAGCGACATCACGCCCCAGCAATGCCCCAACCTGCGGTTCATGACAGTGGGCGGGGAGAAAATCACACAGAAAGTCCTCGACACCTGGGCCTCAGCACCGAACGTGGCTATCGTGAACGC
CTATGGTCCCACGGAGGTAACGATTGGCTGTACATTTGCGCATATCAACCCGTCCACAAACCTACGCAATATCGGCCCCCCTCTTACAGCTTGCACAGCGCATGTTCTGATCCCTGGGACCATGAAG
TACGCACTTCGGGGCCAGACTGGCGAATTATGTTTCAGTGGTGATCTTGTGGCGAGAGGATATCTGAATCGCCCTGATGCCACTGCCGCAAATTTTATCACTGGCCCCAACGGTGCCAAGATGTACC
GTACCGGTGATATCGGCCGGCTTATGTCTGATGATTCGGTTGAGTACCTTGGACGTGGAGATGACCAAACCAAGATCCGTGGGCAGAGACTAGAACTCGGCGAGGTTTCCGAAGTACTAAGAGCTT
CTTCGCCAGTTGCCGTTGATATTGTGACGACGGTAGCGAAACATCCTGACCTCGGCAAGGTACAGCTGATTACCTTCGTGTCGAGAGCAAAGAAGCGCACTGTCAATGAGGAGGTGCAGTTTCTATT
CTCTGATTTCGGTACCCTAGGCCAAGAACTGCGCGATATTTGTGCGAAAAAGCTGCCAGCCTATATGGTCCCAGACCTCATCTTGCCCGTGACAAGTATCCCCGTCGCGGCAATGAGCGGAAAGGC
GGATATGAAGGTATTGCAAAAATTATTTACCGAGCTGCCGTTGCAGGTTGTTCTTCAGGGGAATAATGCTATTGCAAACGGAACTGGCTCCGAACGGCCGCTGAACCCAGACGAGATAGCCGTCGTC
GGCGAGATTTGCCAAGTTATCTCTGCTGACTCACACAGCTTTTCGCCTATGACGAACATCTTTGAAATTGGGATTGACAGTCTCAGTGCAATTGGGCTCTCGGTTAGACTACGTGGAATTGGATATGC
GGCGAGTGTGGCGGCAATTATGGCCAACCCGGTTGTCGAGCAGCTTGCTAGGTTGCCTAGAGCTTCTGAGCATGGAGTTAATGATCATGCAGATTATTTTGCACAGCGGTGCAAGGAGTTGGAAAG
CCAGTATCGAAGCGTTTTTAGGGATGCTGTCGAGGTTGCCGTAGTCAGGCCTTGTCTCCCATTGCAGGAAGGACTGATTGCGCGCTCTATGAACAGTAACAGCGGCGATGGCAAGCTCTATGTCAAT
CATGTCATTCTGCAGCTGAACAAGGATGTTGACACCAGAAAGCTAAAAAGCTCTTGGGAGGATGTGGCGAAGGAGAACGAGATATTGAGGACGGCGTTTGCGCCACTAGAGAAGGAGATTGCGCAG
GTTGTCCTGTCCAATGCTTCTTATCAAATGCAATGGACAGAGGGGAAGTATGAAGATTTGGATGAAGCCATTCAAGCACGAAATGAAAAACAAGGACAGATCTCGAGACTCATTATTGCGGATTTATC
AACTGTTCCTCCGGCACGGTTCCATCTTGCCAGCTCGGCGAGTGGGAAGCCGCTGGCTTTGTTTATTTCGATACATCACGGGCTCTATGATGGCGAATCCTTTGCGATGATGCTGGACGAGGTTGC
GGCACGGTATGAAGGGAGGAAGGTCGGGGAGAGAGGCTCACCGTCTGTCTTTTTGAGGCATGTTTGTTCCCAGGACACAGAGGAGGCGAAGAGGCACTGGATGCAGCAACTTTCTGGATGCACCC
CTACAATATTCCGAGCCAACGGAAATGCAATCAAAAACACCATTTCCATTCGTCGAACTGGCAACGCCAAACTCTCGGACTTAGAAACCCGCTCGTCAACACTCCAGACGACAGTCCCGAACCTGCT
GCAGGCAGTCTTTGCTCTCTTGTTAGCGGACCACACCGGTATTTTCGATGTAACCTATGGCCTTGTCCTGTCAGGACGCACTATATCCGCCCCAGGGGCGGACTCGGTACTTTTACCGGCTATCACC
ACTATCCCTGGGCGTCTCAACATGAGCGACTTGAAAACCGTTAATGACGTGGTGAAGGTTGTTCAGAGAGCTACCGCGAGATCTTTAGACTTCCAGCATACACCCCTCCGCAAGATCCAGCAGTGGC
TGAAGTCCGAAGCCCCATTGTTCGATTGTCTGTTCTCGTATATCCGCGCCACTGCTCCACCTGGCCATAACCTCTGGGCAGAATTGGACAGCCACATGCCGTCTGAGTACCCACTTGCCCTTGAAGT
GCAAGCCGATAATGCGGCTAATACACTCAAATTGGAGTGCATCTTCTCCTCAGACTTTGGACCACGGCAAGTTGGTGAGGAATTCCTTGAGAAGATGGACGCTGTTATTTCTGAAGTTGTCTCTGGTT
CAAGTTTGCCTTTGGAGAACTTCAACACCGTGCGATCAGCTACCTCTGCGTCTCATGGTGCATCCGTACAATGGGACGAGTCAAGCTGGACTTCGTCAGAAAGCAGGATACGAGAAATAACGGCAAC
GTTCTGTGGACTGAATGTCGAGGCTGTATCAAAAGGCGCTTCATTCTTTAGTCTTGGCATAGACTCAGTCACTGCTCTTCAGTTTGCCAGAAGGCTGCGTGATGAGGGATTCAAGGTTTCATCATCGG
AAATTATGCGTTTTTCCTGTGTTGGGTCGTTGACAGGACATATTGAATCTTCTTCCGCGCTGCAAACTAATGGTATTGGGAAAACGGAGACGGCTATCTCAATCGAAACCTATGCAAAGCATATTCCTC

TGCTTGGAAAGAATGACTCCATCACGTCGTTGTTCGAATGTACACCACTGCAATCAGGAATGATCACGCAAACGATCAGCTCTGGAGGGAAGGTGTATATTAATCCTCATCCAATCCGTCTCAGGGAC
AACGTGAAAGTGGAGAAGCTCAAGGAGGCCCTGCGTCACGTTGTTCAGGCTAACGAAATTCTGCGCACGTCTTTTCACCTCATCCCGGATCTAGGAGAGAGCTGGATTGGTGCCGTGCATGAAGAG
CCTAAGTTTGAGTGGAGTGAGATCAATATGCCATCAGGCGCCAATGCTCTGTCTGAGGTTATGAACCTCTATACGTTCTGTGAAGAGGCATCTTTTGAGAGGCCACCGATTCGTTCTGGTCTTGTCAA
CCAGCCAGGCTATAGGATCCTCATAGCTGTCCTACATCATTCCCTTTACGACGGTGCCTACCTACCGTTCGTCTTCGAGGACCTTGCAACAATCTATGCAGGAGGAACCCTTGCTCAACGACCACAG
TTCTCAGAAATGGTCCCCTACATGCTCTCAGGGCAAGACGAATCCTGTGCCTTCTGGGTGGACAGGCTCAGAGACTATGTACCTGTTGAGATCTGTCCACTACCCAAGCTCGACAGCACTCCAAATA
TGCTAACCGCGAAAAATCGCATTCCCCTACCCCTCCACTCAATCACAGAATCCTGCAAGACCATGTCCGTCACAATCCAAACTGTCTCCCTGCTTTCCTACGCAAAGGCCTACGCGCACCTTCTCGG
AACCCGTGACGTCGTGTTCGGTCAAGTCTTAGCCGGCCGAACACTGCCTCACCCTGAGGCAGACCGTACATTAGGACCATTATTCAACACTGTCGCGCAACGAATAACGCTTGATCCCACATTTATG
AGTAACTGCGCTCTTGCGCAACGGCTCCAAAGGGATGGCGTCGAAGCACAAAGACATCAGCATGCACCGCTGAGAATTATTCAGAATACGCTGCGTCAAGAGGGAAATCTTGACTCGAAGCAGTTG
TTTGATGCACTCTTTGTGTTCCAGAAGAGTGCCGCGCTCTCTCAGGGTATCCTGAACGAGCAGGAGATTTGGACATCATATGAAGATGAGGATTTCGTCGTTGACGCGGAATATAAACTCAATGTCGA
GGTTGACCATTCTCACGACGGGGTCATTGTTAGGGCTACGGCAAATGGTGCCTACCTTAACCAGAAAATGCTTGAGAGTTTCCTCAGTCAATACGTCGAGGTTTTCTGTGATGTCGTCGAGCATCCC
GCAAGATGTGTCACGGCGGTACCGCAGGGACTGGGAGGACTGCCGTTAGAACGGGCGAGTTCGAGATTCGGGCAACCGGTGACTGACGGCTCGAAGCCGGATTCTGCGCCCTCTACGCCTGTGC
CAGTGCATGAGGAAACCATTCGATCTGTTCTAGCAGACGTCGTGGGAATCTCTACTGACGATATCAAGCCAACCACGAGCATTTTCAACCTCGGGCTTGACTCTCTTTCAGCTATCAGGCTCGCTTCT
CTTTGCCGCACAAAGGGTTTGAAAGTCAGTGTTGGGGATATTCTTCAGGGGAATACGCTGCGCGGTATAAGTACGCGTGTGCAGTTAGAATCCGAAGTCTCTACCACGACAAACGGGCACAACCCA
GTTCATGGAACTTCGTCCTTGATTAAGGATTATCCTCACGTCGAACAAACCGTCATCTCTAATCTGCATCTCAGCAAAGAAGAGATCGAAACAATTATCCCCGTTCTCCCTGGCCAATTTCACCACCTG
GTTGGCTGGCTCAAGTCAGACCGGAAACTGTTCGAAGCACCCTGGGCTTTTGTCGCTCGTGATGATAAGCGTATCAATGCCGATAAGCTCCAGTGTGCGTGGGCCGATCTTCGCAAGCGACATCCT
GTGCTACGGACGGCCTTTGCAGCTACCTCGGACTCAGAGGCTGTGCAGATTGTTCTGAGAACACCTGCAGAAAACTCGGATGCATTCAGAGTCATCGAATCTGCGGACAATATTGCCGACCTAGCC
AGAGCGCACGCCCGCGAGGAAGCGTTGCACCCATCCTCCCTCTCTTCCCCGCCTGTGCGATTACGGCACCTCAAGGCTGCGGATAGGGATGGCATCCTGCTTATTATCCATGCCTCTCTGTACGAT
GCATGGAGCATTCCGATGCTTGTCTCTGAACTTGGTAAACTCTACGACGATCAGCCAACAGATTTTACCACAGCCCCAGACTTCCCTGCCCTCGTGGACTTTTCACTTCGTGCTCTCTCCAACCTCGA
TGTCAACGAGAAGGATTACTGGACCTCGACTCTTAAGCCCGCTACACCGACTCTTGTCAGGAGCGCCGGTACGCAGAAATCAATTGCAAACAACGAACAGTTGTTCGTTGGCGAATGGGAAAGAGT
CTCCAATCTATCTACGATGGAGAAAATATGTCGATCCGCGGGATTCAGCCTTCAGACGATCATCCTCCTCGCTGTTGCGCGCTGCCTTGCGAGATCCACTGGCGTTGAGAGCCCCGTCATGGGACT
CTATCAGAATGGGCGCTTGGCTGCGTTCGATGGAATTGAAAGGGTCCCGGGACCCTGTTTGAACGTCAATCCGTTTGTTGTGGAGGATGTGTTAACCAGTCTGGGCAATGAAGAGAAAGAGTGTGT
CCTCAAGCAGGCGAGAAACATTCAGAGATCCCTCGCAGAACGGGTACCGTATGAGCAGAGTTCGCTTCGTAAAGTCCTAACATGGCTTAACCCGGAAAATGGAGAGGTAACACCGCTATTCAATATG
TGGGTTAATCTGCTTTGGATGCAGGACAGCACTTCCTCCACCCCCAGCCAGCAAGCCAATGACGAGAAAAGTGAAGCCGGGTTCTTCAAACCCCTTCGCATCGGCGTTCCGACAGACTTCATTCCTT
CTAAGCCGCTACCCCCATCATCTACATCGATAGATAGCCTCGATACATCTTACTTGCCAGACCAGAATATCTTCCTTGATATTGGTCCCGATCCAGCCACGGATAGTATTGGCTTTGGGGTCCGTGTT
GAGGGTGGCTTGCTGGCAGAGGGTGAGGTCAAAGAGCTTGTGGACGCGATTGCCGCTGAGATTGAAAGAGCTGTTGCTTGCTTAAAAGCTCACGTGCATCACCATCACCATCACTAA

#### pHis$_{14}$-SidC NRPS (amino acid)

MASHHHHHHHTAMGKRKLAGSFRRYVATGLSRVSGAFRSRSASKERQSRGSGSLVERDTLAKDQGQGVC
LQPAQVGAVPVTKVPALDTIEAYAGAFDKPIDGAGTCQRTSLVCSASSAVDGDVIDGLVQGYARFIAGLTGLD
DIAFYSTRHEPFALEKSTSIIIASAASNELVCREVDTNHENDVQFYAELGRVEPENGRREFQPNAFTLFIEPDSN
RKKKNVLHLSFAYPRRLIPDAAVEQLLRTLLLHICESSPLKSPSTSQPELSILNFPPSMIPPTAQANGVENSTTN
PHLLHSAFENWARKNPHFIALDFIHSLSSKTHRSEHSIITYAALDSAATNLAFHIRSLLRDSRKHGQIIPVHMPTS
PELYISYLAVLKAGHAFCPIPQDVPARRIQEILSDIDAPIVLGTSSKPPISAESSRSTSTWVNVTEVSKWRQMCG
EQPADYSRPSLDHITIEQNQTAYLLFTSGSTGKPKGVQISHLAASCSISSHATAIPLPGESPGNFRWFQFASPS
FDPSLMEIFVTLSTGGTLCSADRRLTLANLEATINESRATVMMATPSLATLLRPDRLETLEALWSMGEKLNRTV
IDNFALDNVMNGDAETRRPRTLVNAYGPTEGAINCTYVAPFKRHMRGSIIGRPLPTCAMFILSPDSQVPVLVPT
GTVGELAIGGPQVSKGYLNLPEVTARVFIRSKEFGPLYRTGDKARIVWDESGHQVIEYLGRIRTDQVKINGRRV
ELGEIESVVAAVEGVREAVAVVVKRDSKSNGGEQIVACLVVDAADGEGREKIAQQAKKNAAQHLASFMCPTT
YTFFDVLPRSSSGKVDRKALAVQLQEKPDVPIKNGLSEESAECWRHSDEAASSVQQLVIRLVAETGDVSDSAI
KPGTELYSVGIDSLGAMRFLQKLRDNGVHGLSVGEVLQTHTCQRLVSLVQSMLTNQNGEPNGILQKGSITND
LQLRLQSFGRRYRMFCAESLDVPADTIHEVLPTTATQSGMLTSFLRSSAEQSYEKPTYIYHTVLPLEPRTNIEK
LKKAWFDVISNYDSFRTVFCMVDDELAPFAQCILTAEGVSSRDWNVYTSPNECSAENDIIDHALRSAEESITLR
RPPWKLSLVQSSTKSIMILSMFHGIFDGGSLQLLLQDVSSAYSGNTLPQRTSLTHIVKHHFQADHTSTSKFWR
EYLQGYSLLPFPSLTPHRAPAQKSTGCAEVTSQLSYGALKKLSKSIGSTPLSVLQAAWGAVVLSYTATPDQDV
VLGSVMSGRLDPDSKDCIGPTFTITPSRISVQQLKGSGSLTNQSVVHYLSSSNAKALSHLQPQLGSVTVNGKV
PYDTVLAYQDFDTDMKPSQTWSSVQHPAMANDFSVMIEVVPNPDSTLTLRASFDTKLDSTGAQIMLKQMDDI
VSYILNHPDSSFEDAPLQASLALKSKANPSPITAPEVSEGALLQSQFEDHALSHPNDPALVFKQDLDNDDHPG
NITWTYAQLNAMAEALAEHLLQVCGDLRDASVPICIEKSPPLYVAILGILKAGGAWCPIDTLSPAQRRHDLIART
SAGILLVSGLDTPQPQNAVPAGVRVIDVSKFIQNVSSDNTPQSSRHRATPRNTAYLIWTSGTTGAPKGVPITHS
AAVSSMRSLQTDIPGNEDGSPIRCLQFSQPTFDVSIQDLFYTWGLGGALISGTREIMLESFPKLANITKATHAHL
TPAFAAGVARKSCKTLKVVTMIGEKLTQSVADDWGTDMRAFNTYGPAEATVVSTIREFGNEHRSVKSANIGW
PMSSVSVFVMSKDKRVLMKNAIGELALGGPQLSPEYLNLKDVTDTKYIWNEDAGQRLYYTGDLVRMLSDGSL
EYITRVDDLVKLGGIRIELSEISFALRGCHELVESVETMILSRKDRPVRVVVAFLCAPKAAGDADEGLLVLDDTG
RDIARAASLQARNVLPENMIPSVYLIVKKIPKTPSAKVDRRALQAAYAAVDIDKWENNVNPEGPGDADEDDAA
TATQIIETVAALVHVESSTITKSNRLRSLGVDSLCATRLAFRLKEAGFGLSVMDVLACTTIQDLVRLAQSTSFSS
SANSASPHNDKFDIVSFNKIWHTLVAAAAKIPEKDAFTTIRATAIQESLLTETMGTYDMYWSNHFFRLDRSVDIP
RLRQAWYAVCQKTETLRTGFIPVAQTEAKNKKQAKDSGFSILQVVYKLPAVDWEAHTYRKEEWSWVLKNRV
RDIMTAHQKNYFCHPPWAVTILEEGTERVMVLTLHHSIHDEPSLKFLMDDVRAAYTYKPPLRTQLTPALALVLP
TPSKYAEAVDFWSLELKPYAALDVPVWPDLTGKRVSPGAAPEYKLISEAMSITSSFPVLEKVAADLGLKSVASII
RAAWAFVSLSYLGLSGTVFAETLSDRVFDPSLESAVGPFISVVPVPFRVEGDTTVRKILAEQHRLSLQSWKHR
HVHARDIRKALKRQRGEPLYPAVFNFHALDESKDRKIASLPGLWHELEDQIGLHVEHPMAMNVFQSPSGNMT
LEASSDSRIFSREHLRLFVRQIDALISEMLLSPDESLSGLVNRLPSDLRSLSNRIVSHEVANSIHQAPTYWLEKF
ADTHPHWTAVEVASSISTNGIEKEAMSYGSLNSAANRVAAYLASFRYKNRVVGVCAGRTLASYPIIIGIFKSGN
TYLPIDESLPADRKAFLLEDAKCPVVFTELGLRNSFAGAPDTCRVECIDDPALQRSLDEMPSTNKDYSSHPDD
VSYLLFTSGSTGKPKGVMVTRANLSSFIESISEFACRIAPDTLKLGGTGRYLAQANRAFDPHLLEMFFPWRHG
MATVTAPRPMILDDIGTTLSKWSITHASFVPSLVDQSDITPQQCPNLRFMTVGGEKITQKVLDTWASAPNVAIV

NAYGPTEVTIGCTFAHINPSTNLRNIGPPLTACTAHVLIPGTMKYALRGQTGELCFSGDLVARGYLNRPDATAA
NFITGPNGAKMYRTGDIGRLMSDDSVEYLGRGDDQTKIRGQRLELGEVSEVLRASSPVAVDIVTTVAKHPDLG
KVQLITFVSRAKKRTVNEEVQFLFSDFGTLGQELRDICAKKLPAYMVPDLILPVTSIPVAAMSGKADMKVLQKL
FTELPLQVVLQGNNAIANGTGSERPLNPDEIAVVGEICQVISADSHSFSPMTNIFEIGIDSLSAIGLSVRLRGIGY
AASVAAIMANPVVEQLARLPRASEHGVNDHADYFAQRCKELESQYRSVFRDAVEVAVVRPCLPLQEGLIARS
MNSNSGDGKLYVNHVILQLNKDVDTRKLKSSWEDVAKENEILRTAFAPLEKEIAQVVLSNASYQMQWTEGKY
EDLDEAIQARNEKQGQISRLIIADLSTVPPARFHLASSASGKPLALFISIHHGLYDGESFAMMLDEVAARYEGRK
VGERGSPSVFLRHVCSQDTEEAKRHWMQQLSGCTPTIFRANGNAIKNTISIRRTGNAKLSDLETRSSTLQTTV
PNLLQAVFALLLADHTGIFDVTYGLVLSGRTISAPGADSVLLPAITTIPGRLNMSDLKTVNDVVKVVQRATARSL
DFQHTPLRKIQQWLKSEAPLFDCLFSYIRATAPPGHNLWAELDSHMPSEYPLALEVQADNAANTLKLECIFSS
DFGPRQVGEEFLEKMDAVISEVVSGSSLPLENFNTVRSATSASHGASVQWDESSWTSSESRIREITATFCGL
NVEAVSKGASFFSLGIDSVTALQFARRLRDEGFKVSSSEIMRFSCVGSLTGHIESSSALQTNGIGKTETAISIET
YAKHIPLLGKNDSITSLFECTPLQSGMITQTISSGGKVYINPHPIRLRDNVKVEKLKEALRHVVQANEILRTSFHLI
PDLGESWIGAVHEEPKFEWSEINMPSGANALSEVMNLYTFCEEASFERPPIRSGLVNQPGYRILIAVLHHSLYD
GAYLPFVFEDLATIYAGGTLAQRPQFSEMVPYMLSGQDESCAFWVDRLRDYVPVEICPLPKLDSTPNMLTAK
NRIPLPLHSITESCKTMSVTIQTVSLLSYAKAYAHLLGTRDVVFGQVLAGRTLPHPEADRTLGPLFNTVAQRITL
DPTFMSNCALAQRLQRDGVEAQRHQHAPLRIIQNTLRQEGNLDSKQLFDALFVFQKSAALSQGILNEQEIWTS
YEDEDFVVDAEYKLNVEVDHSHDGVIVRATANGAYLNQKMLESFLSQYVEVFCDVVEHPARCVTAVPQGLG
GLPLERASSRFGQPVTDGSKPDSAPSTPVPVHEETIRSVLADVVGISTDDIKPTTSIFNLGLDSLSAIRLASLCR
TKGLKVSVGDILQGNTLRGISTRVQLESEVSTTTNGHNPVHGTSSLIKDYPHVEQTVISNLHLSKEEIETIIPVLP
GQFHHLVGWLKSDRKLFEAPWAFVARDDKRINADKLQCAWADLRKRHPVLRTAFAATSDSEAVQIVLRTPAE
NSDAFRVIESADNIADLARAHAREEALHPSSLSSPPVRLRHLKAADRDGILLIIHASLYDAWSIPMLVSELGKLY
DDQPTDFTTAPDFPALVDFSLRALSNLDVNEKDYWTSTLKPATPTLVRSAGTQKSIANNEQLFVGEWERVSNL
STMEKICRSAGFSLQTIILLAVARCLARSTGVESPVMGLYQNGRLAAFDGIERVPGPCLNVNPFVVEDVLTSLG
NEEKECVLKQARNIQRSLAERVPYEQSSLRKVLTWLNPENGEVTPLFNMWVNLLWMQDSTSSTPSQQANDE
KSEAGFFKPLRIGVPTDFIPSKPLPPSSTSIDSLDTSYLPDQNIFLDIGPDPATDSIGFGVRVEGGLLAEGEVKEL
VDAIAAEIERAVACLKAHVHHHHHH

**Table S1.** Primers used for the cloning of SidC constructs and mutagenic primers to generate point-mutations/truncations.

| **Primer** | **Sequence (5'-3')** |
|---|---|
| pLH-SidC-WT-F1 | TGGCTAGCCATCACCATCACCATCACCATCACACTGCAATGGGGAAACGTAAGCTGGCTG |
| SidC-WT-R1 | GATATTCCCTCCAGAATTTGGATGTGGAAGTGTG |
| SidC-WT-F2 | CACATCGTCAAACACCACTTCCAGGCAG |
| SidC-WT-R2 | CTGCGACTTTCTCTAAAACTGGGAAGGACG |
| SidC-WT-F3 | GTCAATAACCTCGTCCTTCCCAGTTTTAGAGAAAGTCG |
| SidC-WT-R3 | CCGCAATAATGAGTCTCGAGATCTGTCCTTG |
| SidC-WT-F4 | GAAATGAAAAACAAGGACAGATCTCGAGACTCATTATTGCGG |
| pLH-SidC-WT-R4 | AATTAGTGATGGTGATGGTGATGCACGTGAGCTTTTAAGCAAGCAACAGCTCTTTCAATC |
| pLH-SidC-$T_1$-F | GCTAGCCATCACCATCACCATCACCATCACAGTGAAGAAAGTGCAGAATGTTGGAGGCAT |
| pLH-SidC-$C_2$-F | GCTAGCCATCACCATCACCATCACCATCACAGTATAACAAACGACTTGCAGTTAAGACTG |
| pLH-SidC-$A_2$-F | gCTAGCCATCACCATCACCATCACCATCACGAAGTCAGCGAAGGAGCACTCC |
| pLH-SidC-$C_3$-F | TAGCCATCACCATCACCATCACCATCACGATCTTGTCAGATTAGCCCAGTCTACTTC |
| pLH-SidC-$T_4$-R | CGCGCTGCAAACTAATGGTATTTAATTTAAATGACAAATTTGTGCGGACCGATGCC |
| pET-SidC-$A_1$-F | TACTTCCAATCCAATGCAGCAATGGGGAAACGTAAGCTGGCTG |
| pET-SidC-$A_1$-R | TTATCCACTTCCAATGTTATTAACTTTCTTCACTGAGGCCATTCTTGATTGG |
| pET-SidC-$T_1$-F | TACTTCCAATCCAATGCAAGTGAAGAAAGTGCAGAATGTTGGAGGC |
| pET-SidC-$T_1$-R | TTATCCACTTCCAATGTTATTACTCGCCATTTTGATTCGTCAGCATGCTC |
| pET-SidC-$C_2$-F | TACTTCCAATCCAATGCAAGTATAACAAACGACTTGCAGTTAAGACTGCAGTC |
| pET-SidC-$A_2$-F | TACTTCCAATCCAATGCAAGGGAGCACTCCTACAATCACAGTTCG |
| pET-SidC-$A_2$-R | TTATCCACTTCCAATGTTATTACTCAGGGTTAACGTTGTTCTCCCACTTGTCAATATC |
| pET-SidC-$T_2$-F | TACTTCCAATCCAATGCAGGAGATGCAGATGAGGATGATGCTGCC |
| pET-SidC-$T_2$-R | TTATCCACTTCCAATGTTATTAACTGAATGAAGTAGACTGGGCTAATCTGACAAGATC |
| pET-SidC-$C_3$-F | TACTTCCAATCCAATGCAGATCTTGTCAGATTAGCCCAGTCTACTTCATTC |
| pET-SidC-$T_3$-R | TTATCCACTTCCAATGTTATTATCCATGCTCAGAAGCTCTAGGCAACC |

| | |
|---|---|
| pET-SidC-$C_4$-F | TACTTCCAATCCAATGCAGGAGTTAATGATCATGCAGATTATTTTGCACAGCG |
| pET-SidC-$T_4$-F | TACTTCCAATCCAATGCATCTGCGTCTCATGGTGCATCCGTACAATG |
| pET-SidC-$T_4$-R | TTATCCACTTCCAATGTTATTACTCCGTTTTCCCAATACCATTAGTTTGCAGCG |
| pET-SidC-$C_5$-F | TACTTCCAATCCAATGCAACGGCTATCTCAATCGAAACCTATGCAAA |
| pET-SidC-$T_5$-F | TACTTCCAATCCAATGCAGATTCTGCGCCCTCTACGCCTGTG |
| pET-SidC-$T_5$-R | CCGTTATCCACTTCCAATGTTATTAGGTAGAGACTTCGGATTCTAAC |
| pET-SidC-$C_T$-R | CCGTTATCCACTTCCAATGTTATTttaAGCTTTTAAGCAAGCAACAGCTCTT |

**Table S2**. Measured and calculated (in brackets) masses of *apo*-, *holo*- and haOrn-bound species detected by intact protein MS. All calculated values are average protein masses.

| Protein | *apo-* | *holo-* | *holo*-L-haOrn | *holo*-(2 x L-haOrn) | *holo*-($Gly_3$-L-haOrn) |
|---|---|---|---|---|---|
| **SidC $C_3A_3T_3$** | 126,449 Da (126,452 Da) | 126,790 Da (126,792 Da) | 126,963 Da (126,964 Da) | - | 127,136 Da (127,133 Da) |
| **SidC $C_3A_3T_3C_4T_4$** | 188,652 Da (188,656 Da) | 189,331 Da (189,336 Da) | 189,506 Da (189,508 Da) | 189,680 Da (189,682 Da) | - |
| **SidC $C_3A_3T_3^0C_4T_4$** | 187,571 Da (187,574 Da) | 187,911 Da (187,913 Da) | 188,085 Da (188,086 Da) | - | - |
| **SidC $C_4T_4$** | 60,270 Da (60,271 Da) | 60,611 Da (60,611 Da) | 60,785 Da (60,784 Da) | - | - |
| **SidC $T_4$** | 9,911 Da (9,912 Da) | 10,251 Da (10,252 Da) | - | - | - |
| **SidC $C_5T_5C_T$** | 117,484 Da (117,492 Da) | 117,821 Da (117,832 Da) | 117,994 Da (118,004 Da) | - | - |
| **SidC $T_5C_T$** | 64,641 Da (64,644 Da) | 64,980 Da (64,984 Da) | - | - | - |
| **SidC $C_2A_2T_2$** | 163,865 Da (163,866 Da) | 164,205 Da (164,206 Da) | - | - | - |
| **SidC $C_2^0A_2T_2$** | 163,799 Da (163,800 Da) | 164, 138 Da (164,140 Da) | - | - | - |

**Table S3**. Measured and calculated (in brackets) masses of Gly-bound species detected by intact protein MS. All calculated values are average protein masses.

| Protein | *holo*-Gly | *holo*-$Gly_2$ | *holo*-$Gly_3$ | *holo*-$Gly_4$ | *holo*-$Gly_5$ |
|---|---|---|---|---|---|
| **SidC $C_2A_2T_2$** | 164,262 Da (164,262 Da) | - | 164,378 Da (164,377 Da) | - | 164,490 Da (164,491 Da) |
| **SidC $C_2^0A_2T_2$** | - | - | 164,307 Da (164,311 Da) | - | 164,421 Da (164,425 Da) |
